# De novo Rubisco design with protein language models

**DOI:** 10.64898/2026.09.04.749267

**Authors:** Alexander J. Kehl, Simon K. S. Chu, Jose Henrique Pereira, Jennifer Lee, Renee Z. Wang, Michael Gigl, Paul D. Adams, Patrick M. Shih, Justin B. Siegel

## Abstract

Ribulose-1,5-bisphosphate carboxylase/oxygenase (‘Rubisco’) fixes the majority of carbon dioxide globally but is challenged with low specificity for CO_2_ versus O_2_ and low catalytic efficiencies. Traditional engineering efforts have remained difficult because folding, assembly, specificity, and catalysis are tightly coupled, hampering efforts to explore sequence space. Therefore, we leveraged recent advances in protein large language models (PLMs) to generate sequences beyond those observed in nature, using both ProGen-2 that was fine-tuned on a limited dataset of non-Form I Rubiscos and an ESM-2 discriminator. With this approach, we generated 5.6 million novel Rubisco-like sequences and identified 21 highly diverse candidates predicted to be active that occupy regions of Rubisco phylogenetic space not previously observed in nature. Six designs were soluble in *Escherichia coli*, and five were shown to produce quantifiable 3PGA. One design produced an apparent CO_2_/O_2_ specificity estimate beyond the range of the natural representative Rubiscos assayed. We also solved the crystal structure of one de novo design that reproduced the predicted dimer and active-site geometry with sub-angstrom Cα agreement. Sequence-only generation followed by independent structural filtering therefore recovered soluble, active Rubiscos from regions of sequence space that are not represented in genomic databases. Together, these results establish a scalable strategy for accessing previously unexplored Rubisco sequence space, providing a broadly accessible path toward generating de novo Rubiscos that have activity and specificity parameters needed to address longstanding limitations in biological carbon fixation.

## Introduction

Rubisco is the principal carboxylating enzyme in autotrophic metabolism and the major biological entry point for inorganic carbon (*1–3*). The enzyme arose early and diversified across all domains of life (*4*, *5*). The Rubisco superfamily comprises several phylogenetically and structurally distinct lineages—Forms I, II, II/III, III-Like, IIIA, IIIB, IIIC—whereas the related Form IV Rubisco-like proteins (RLPs) lack canonical RuBP carboxylase activity (*4*, *5*); we note that the naming of these clades is in flux and refer to the naming as discussed in Kehl et al. 2026. Its recognized forms differ in sequence, metabolic context, oligomeric state, and accessory requirements. Large subunits assemble to form L_2_ dimers, which then assemble higher-order homo-oligomers, or assemble with the small subunit to form L_8_S_8_ complexes (*6–10*).

In contrast to this phylogenetic and oligomeric diversity, native rubiscos display relatively slow rates of carboxylation and a competing oxygenation reaction. In organisms which rely on rubisco for carbon fixation, this sets limits on growth, motivating engineering strategies to increase ‘specificity’ for CO_2_ over O_2_ (*11*). Two parameters describe this limitation: the carboxylation turnover number, k_cat,C_, the number of CO_2_ molecules fixed per active site each second, and the CO_2_/O_2_ specificity factor, S_C/O_, which determines how carboxylation and oxygenation are partitioned at a given CO_2_/O_2_ ratio.(*12*, *13*) Directed evolution and structure-guided mutagenesis have improved k_cat,C_, S_C/O_, or both in individual Rubiscos (*14–18*), including through selection for altered oxygen response (*19*). Broad engineering strategies remain limited by rugged sequence– function landscapes (*20–22*) and by chaperone-dependent folding and assembly in heterologous hosts (*23*), which are needed for high-throughput screens of generated mutants. Even modest sequence changes can disrupt several interdependent properties, leaving sparse islands of function in the surrounding sequence space. Structure-guided approaches additionally depend on explicit mechanistic models, or on representative conformational states, for a large, dynamic, interfacial active site. These requirements have limited their use for broad searches of the sequence landscape (*8*).

Protein language models (PLMs) offer a complementary design strategy to broadly search the sequence landscape, by learning sequence constraints directly from natural proteins (*24*). Models trained at evolutionary scale encode contextual residue relationships and can be specialized by family-level fine-tuning (*25*, *26*). This approach has generated functional lysozymes (*27*) and CRISPR-Cas proteins (*28*). Unlike pairwise coevolutionary models (*29–31*), PLMs can represent higher-order dependencies without requiring a fixed structural template. They can therefore generate combinations extending beyond any single observed sequence or starting structure, enabling broad searches of sparsely sampled protein sequence space. Downstream structure prediction then provides an independent test of fold and assembly quality.

Here, we combined existing methods, using a ProGen-2 generator fine-tuned on natural Rubiscos excluding Forms I and RLPs with an ESM-2 discriminator, sequence clustering, and structural filtering (**Figure 1**) (*25*, *26*). We refer to this unified generative–discriminative workflow as RubisGen. We evaluated 21 designs by expression, protein thermal-shift analysis, endpoint LC– MS, and crystallography and compared them with a phylogenetically diverse panel of natural representatives. The resulting proteins show that sequence-only generation followed by independent structural filtering can recover soluble, thermally stable, 3PGA-forming Rubiscos that retain the canonical dimeric architecture while exploring new sequence and functional space beyond what was previously available from genomic databases.

**Figure 1.**
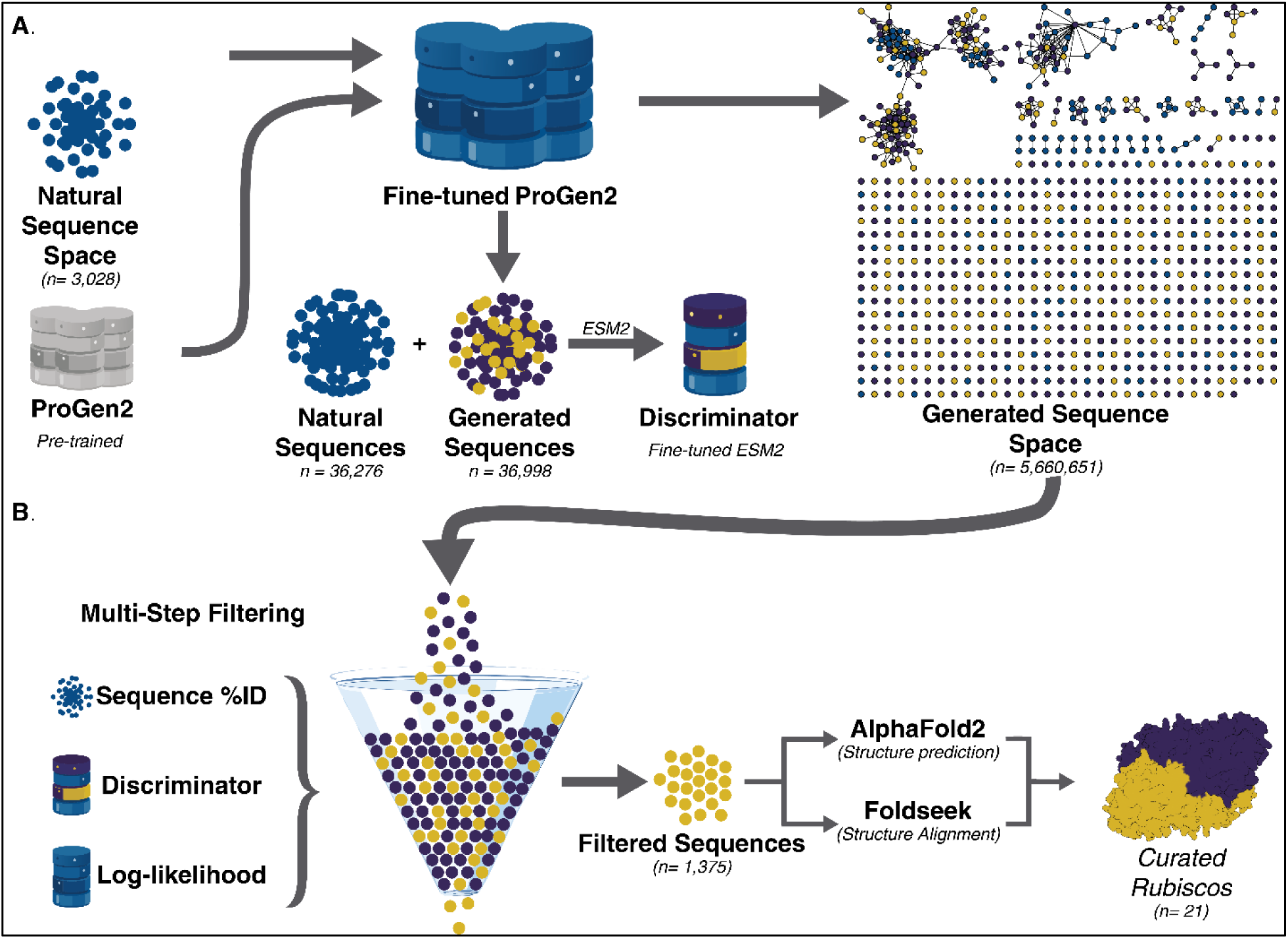
RubisGen identifies Rubisco candidates for experimental validation. **(A)** ProGen-2 was fine-tuned on 3,028 stringently curated full-length natural Rubiscos excluding Forms I and RLPs and sampled to generate approximately 5.6 million candidates. Candidates were scored independently by an ESM-2 discriminator reporting p(generated). **(B)** Natural-sequence identity, p(generated) (Discriminator), Log-likelihood (generator loss), MMseqs2 clustering, AlphaFold-multimer prediction, Foldseek comparison, and active-site inspection reduced the candidate set to 21 designs for experimental validation.

## Results

### RubisGen identifies Rubiscos across diverse sequence lineages

We fine-tuned ProGen-2 on 3,028 natural Rubisco sequences after excluding Forms I and RLPs, focusing generation on bona fide large-subunit-only Rubiscos and avoiding the additional assembly requirements of Form I systems (*5*, *8*). This generator set was stringently restricted to coherent full-length proteins, which we set by amino acid length (361–500 residues). Candidate scoring used a separately trained ESM-2 discriminator containing 36,276 natural sequences after exclusion of RLPs and 36,998 generated sequences. The discriminator retained a broader 200– 500-residue set, including fragments, because its natural-versus-generated embedding classification did not require every input to satisfy the full-length generator criteria. Nucleus sampling at top-*p* values of 0.5, 0.75, and 1.0 produced approximately 5.6 million unique sequences; lower top-*p* values restrict each sampled residue to a smaller set of high-probability model outputs. Generator loss measured agreement with the learned sequence distribution, whereas the ESM-2 discriminator returned p(generated); lower values indicated a more natural-like classification.

Sequence scoring and MMseqs2 clustering at 80% identity (*32*) reduced the generated set to 1,375 candidates for AlphaFold-multimer prediction (*33*) and Foldseek search against the PDB (*34*). The unified selection funnel combined low p(generated) or generator loss with natural-sequence identity, high predicted confidence, removal of Form I and RLPs structural matches, and manual inspection of active-site completeness; candidates also underwent ProteinMPNN and spatial aggregation propensity filters (*35*, *36*). The 21 selected designs had p(generated) values of 0.000795–0.351, mean pLDDT values of 90.33–95.19, and top-hit Foldseek identities of 0.423– 0.694. All five distinct top PDB hits were Rubisco structures (PDB 2D69, 3WQP, 5MAC, 6HUN, and 8DHT; **Table S1**), confirming that the structural screen retained the intended fold rather than generic similarity to unrelated PDB entries.

In the sequence-similarity network generated at an alignment-score threshold of 70, RubisGen sequences occupied multiple regions rather than forming a single separate cluster (**Figure 2A**). The 1,375 structurally evaluated designs combined high mean AlphaFold pLDDT with a broad range of Foldseek identities (**Figure 2B**), and their phylogenetic placements covered several lineages, including sparsely sampled deep branches (**Figure 2C**). Thus, RubisGen retained candidates across multiple sequence and structural neighborhoods rather than converging on one natural template.

We also asked how far the generated sequences had moved from the natural collection they were compared against. Of the 1,375 structurally evaluated designs, 1,243 had a match in the original sequence database and 132 had none; the designs with a match shared a median of 59.73% identity with their closest natural sequence (interquartile range 54.02–72.09%), and the designs without a match were kept as a separate group rather than being recorded as zero identity (**Figure S1**). A two-dimensional map of the sequence-similarity-network set, which contains 987 generated and 639 natural representative sequences, placed the generated sequences across several natural neighborhoods rather than in one isolated region, in agreement with the network in Figure 2A (**Figure S2**). For the five designs that later produced quantifiable 3PGA, identity to the nearest natural sequence was 70.6–80.4%, the longest stretch of consecutive identical residues shared with any distinct natural match was 19–52 residues, and the longest such stretch within each design’s closest match was 16–50 residues (**Figure S3; Tables S11a and S11b**). In a randomization test of the closest match, the identical residues were more clustered than expected by chance for GEN_4, GEN_7 and GEN_10, but not for GEN_8 or GEN_11 (**Table S11b**). Differences from the closest natural match were therefore distributed along the sequence while many positions equivalent to ligand-contact residues were retained.

**Figure 2.**
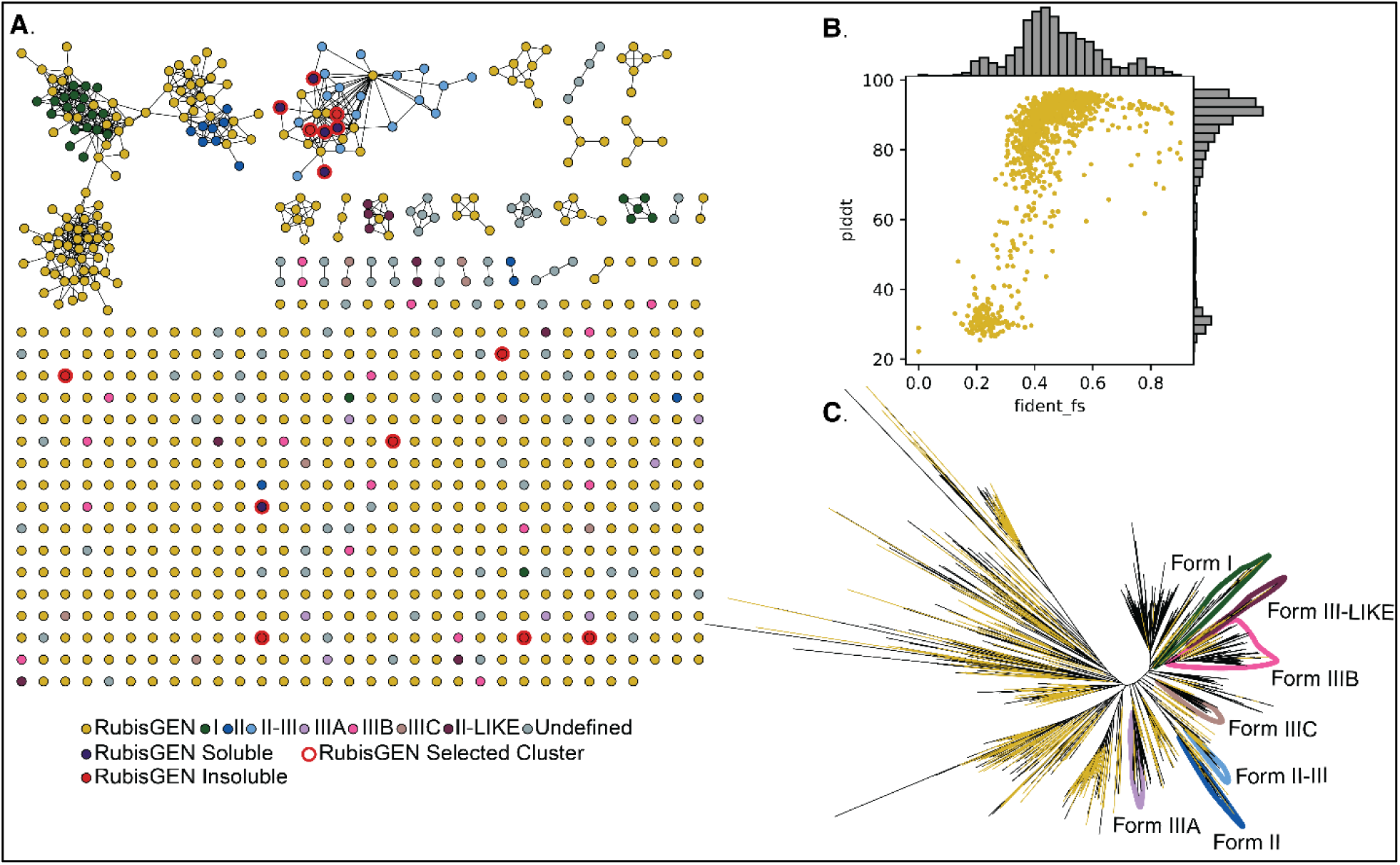
Generated Rubiscos span broad sequence and structural diversity. **(A)** Sequence-similarity network generated at an alignment-score threshold of 70; the representative-node network uses a 40% identity grouping setting. Natural sequences are colored by annotated Rubisco clade: green, Form I; dark blue, Form II; light blue, Form II–III; purple, Form IIIA; pink, Form IIIB; tan, Form IIIC; brown, Form II-like; gray, undefined; generated sequences are gold. RubisGEN Soluble: dark purple, RubisGEN Insoluble: red, RubisGEN Selected Cluster: red ring **(B)** Mean AlphaFold pLDDT versus Foldseek structural identity for 1,375 designs; each point represents one design, and the marginal histograms show the corresponding score distributions. **(C)** Phylogenetic placement of generated sequences (n=987) and wild-type rubiscos (n=639) in an unrooted phylogeny of the rubisco large subunit, RbcL.

### RubisGen-designed enzymes exhibit carboxylation activity within the natural Rubisco range

To evaluate the 21 RubisGen sequences predicted to fold and likely catalyze the Rubisco chemistry we screened the enzymes – in parallel with 19 representative natural homologs for which RubisGen was trained on – using an LCMS based assay in which 3PGA and 2PG could be measured and quantified (**Methods**). Briefly, purified enzyme was incubated with RuBP for 24 hours, after which the reaction was stopped and the 3PGA and 2PG that had accumulated were quantified in a single endpoint measurement. Synthetic genes for the 21 RubisGen designs were synthesized by Twist Bioscience, and plasmids for the 19 natural representatives were taken from the previously described library; corresponding proteins were produced as previously described via heterologous expression in E. coli with subsequent His-tag affinity chromatography (*5*). Of the 21 RubisGen proteins, six produced soluble protein using these methods, and five produced quantifiable 3PGA. Of the representatives used to train RubisGen, 19 were found to produce soluble proteins and 18 produced quantifiable 3PGA.

The natural representative proteins sampled Form II, Form Iβ, Form II–III, Form III-like, and unclassified Rubiscos across the Rubisco superfamily (**Table S2**) (*5*). Apparent 3PGA specific activity spanned more than two orders of magnitude. *Gallionella sp.* was highest at (1.84 ± 0.05) × 10^−3^ µmol 3PGA mg^−1^ min^−1^, followed by *Bacterium* RKZ25299 and *R. faecalis*. *Ca. Woesearchaeota* did not produce quantifiable 3PGA. This phylogenetically broad range of specific activities provided the benchmark against which the generated proteins were compared.

Five RubisGen designs occupied distinct regions of that distribution (**Figure 3C; Tables S3–S4**). GEN_11 and GEN_8 reached the high-activity region at (3.08 ± 0.10) ×10^−4^ and (3.02 ± 0.13) × 10^−4^ µmol 3PGA mg^−1^ min^−1^, respectively. GEN_7 fell in the middle of the distribution, whereas GEN_4 and GEN_10 were near the lower end. GEN_5 did not produce quantifiable 3PGA. Apparent 2PG activity was reportable for 11 natural representatives and for GEN_8 and GEN_11, designs falling within the measured natural representative range. Protein thermal-shift measurements gave melting temperatures from 47.8 to 72.5 °C (**Table S4 and Figure S4**). Compared with the natural representatives with measured values, the RubisGen designs showed a lower T_m_ distribution (median, 54.0 versus 70.3 °C), with partial overlap between the two groups (**Tables S3–S4**). These comparisons show that RubisGen recovered function across the assayed natural activity distribution rather than only at its lower boundary.

Applying the plate-specific negative-control mean + 2 SD criterion, and requiring raw 2PG to be strictly greater than both that threshold and the lowest quantified 2PG standard, yielded apparent CO_2_/O_2_ specificity (S_C/O,app_) estimates for 13 of the 23 proteins that produced quantifiable 3PGA in this assay, hereafter 3PGA-positive (**Figure 3D and Tables S5–S6**). GEN_8 produced the highest estimate in the measured panel (5.62 ± 0.37), extending beyond the natural representative range measured here, and GEN_11 ranked fifth overall (1.51 ± 0.21). GEN_8’s estimate was based on two passing technical replicates; its first replicate fell below the plate-specific threshold and was excluded. These estimates suggest that GEN_8 and GEN_11 may have high true specificity and nominate them for conventional kinetic measurement, but an endpoint estimate of this kind does not establish the conventional kinetic specificity factor, S_C/O_.

**Figure 3.**
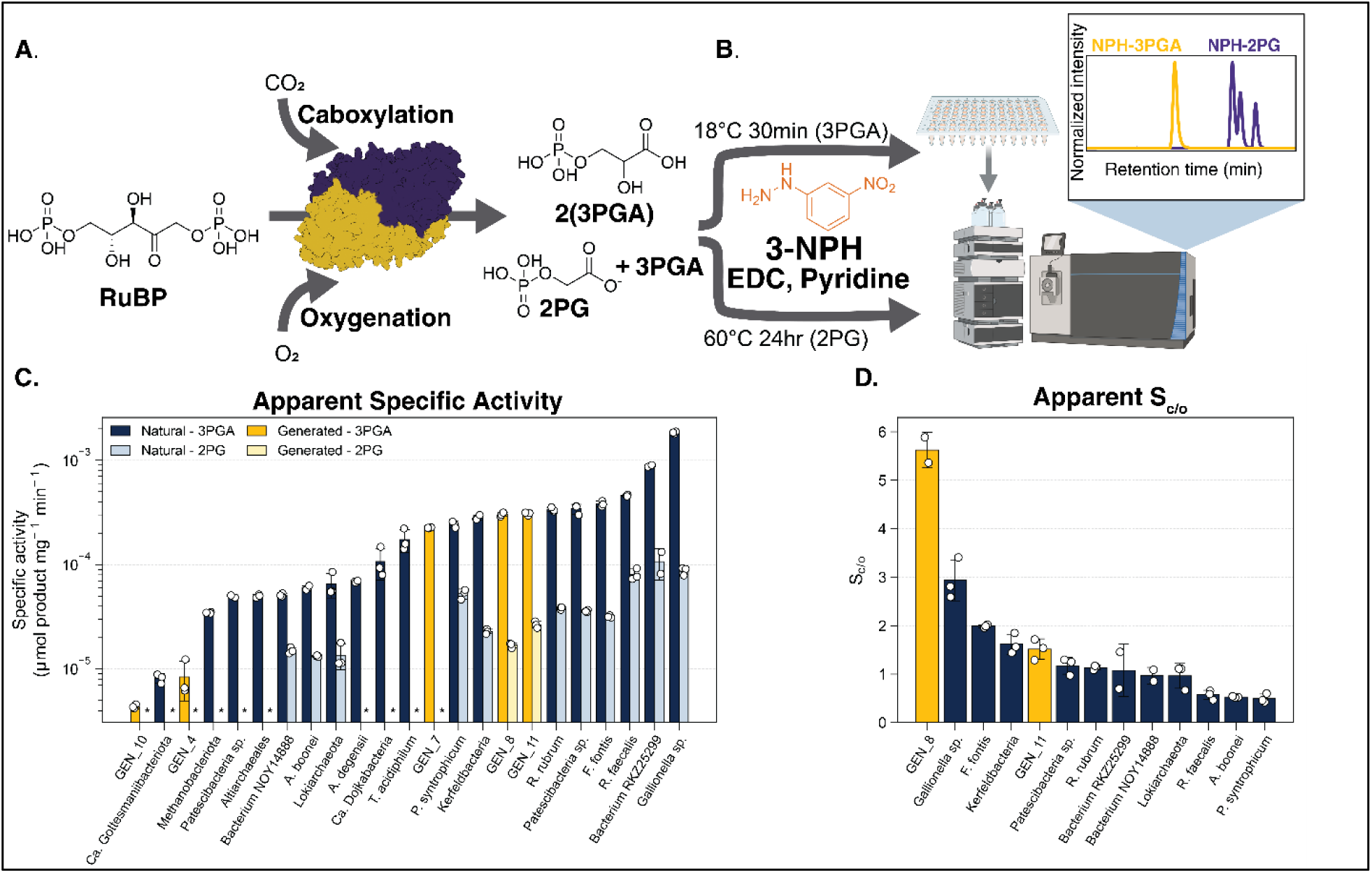
Generated Rubiscos occupy the natural-panel activity distribution and apparent CO_2_/O_2_ specificity (S_C/O,app_) estimates. **(A)** Purified Rubisco was activated for 20 min and then incubated with 1 mM RuBP for 24 hours at 30 °C under 5% CO_2_ and 21% O_2_. **(B)** Products were derivatized with 3-nitrophenylhydrazine in the presence of 1-ethyl-3-(3-dimethylaminopropyl)carbodiimide (EDC) and pyridine and quantified by LC–MS. **(C)** Apparent endpoint 3PGA and apparent 2PG specific activities. Bars show means, error bars show SD, and open circles show technical replicates. 3PGA is shown for all 23 entries; apparent 2PG activity is shown only for the 13 entries with at least two replicates meeting the specificity-reporting criterion. For those entries, all available original 2PG replicates contribute to the activity means, SDs, and points. Missing 2PG bars indicate that 2PG was not reliably detected under the applicable plate-specific reporting criterion and are represented by *. **(D)** Apparent specificity (S_C/O,app_) estimates for entries with at least two technical replicates whose raw 2PG was strictly greater than both the matched plate-control mean + 2 SD and the lowest quantified 2PG standard. The matched plate-control mean was subtracted from passing 2PG concentrations before S_C/O,app_ calculation; 3PGA was not corrected for background because no background 3PGA was detected. Only passing replicates contribute to S_C/O,app_ means, SDs, and points. Natural representatives are blue and generated designs are gold. Panel B was created in BioRender (Vater, A., 2026; https://BioRender.com/j6hjqoy).

### Sequence and structure reveal distributed constraints on function

To test whether the measured range of apparent activity and specificity could be explained by a simple sequence or structural feature, we compared conservation, ligand-contact residues, dimer interfaces, whole-protein similarity, active-site geometry, and static cavity descriptors (**Figure S5; Tables S7–S8**). The canonical Rubisco active-site residues are strongly conserved among natural enzymes, yet no residue was conserved in all 23 proteins that produced 3PGA while differing, or having no structural counterpart, in either of the two purified proteins that produced no quantifiable 3PGA(*37–39*). This was assessed using a sequence alignment (*40*), a structure-based alignment guided by the 9RUB coordinates (*41*), or a whole-protein structural superposition made without reference to the sequence alignment (*42*). The structure-based comparison identified 34 positions identical across all 23 3PGA-positive proteins. However, all canonical ligand-contact residues were conserved in GEN_5 and the natural *Ca. Woesearchaeota* Rubisco, both of which fell below the assay’s detection limit. Their lack of quantifiable 3PGA therefore cannot be explained by the loss of a single conserved ligand-contact residue.

Whole-protein comparisons likewise revealed no gross differences in fold or dimer architecture: median pairwise TM-scores were 0.921 for monomers and 0.932 for dimers, and the closest structural neighbor of each protein without quantifiable 3PGA was itself 3PGA-positive. These comparisons used the structural scoring and alignment methods described in Methods (*42*, *43*). We tested 127 relationships, specified before any statistical result was inspected, between apparent activity or apparent specificity and sequence or static structural features. These features included residues that contact the ligand, internal protein packing, contacts between the two protein subunits (the dimer interface), active-site geometry, overall structural similarity, and access-tunnel properties calculated with CAVER (*44*). We applied the Benjamini–Hochberg false-discovery-rate correction; none of these relationships remained significant after correction except for the signed basic-minus-acidic residue fraction which is a proxy for solubility. Soluble designs had a lower fraction than nonsoluble designs (median 0.02038 versus 0.06378; Cliff’s delta = −0.889; P = 0.000700; q = 0.00980) (*45*). Together, the results argue against a single residue or static structural-nativeness score and instead point to distributed, context-dependent constraints on folding and catalysis (*20*, *22*).

### GEN_10 provides a structural test of a divergent, low-performing design

GEN_10 was selected for structural characterization for three reasons. It had high predicted structural confidence, it produced quantifiable 3PGA near the lower end of the measured distribution, and it occupied an unusually isolated position in sequence space. GEN_10 is a singlet in the sequence-similarity network and is less than 60% identical to every other natural or generated cluster in the dataset. This combination made GEN_10 a stringent test of whether a divergent, low-performing generated sequence retained the predicted fold, dimer interface, and catalytic configuration rather than failing through a gross structural error. We determined a 1.40 Å crystal structure whose refined L_2_ dimer has the canonical Rubisco large-subunit architecture, including the N-terminal domain, TIM barrel, and C-terminal region (*1*, *8*). *R*_work_ and *R*_free_ were 0.149 and 0.170, respectively, and both active sites contain carbamylated catalytic lysine and Mg^+2^ (**Figure 4, Table S9**).

Superposition with the AlphaFold model gave Cα RMSDs of 0.621 Å for the dimer, 0.747 Å for the monomer, and 0.557 Å for the active site. Catalytic-pocket side chains surrounding Mg^+2^ and the RuBP-binding cleft also aligned with the model (**Figure 4C**). Sequence–structure mapping against activated *R. rubrum* Rubisco (PDB 9RUB) showed selective conservation: exact identity among aligned positions was 41.1% for reference residues 1–150, 46.2% for residues 151–420, and 29.2% for the C-terminal region from residue 421 onward, whereas 28 of 35 eligible rigid active-site positions (80.0%) retained the same amino acid identity. Among the 31 structural inliers retained for the local fit, 26 were identical (83.9%); identity across the 475-position full gapped alignment was 39.2% (**Table S10**). The same L_2_ organization and overall fold are retained relative to ligand-free, non-activated *R. rubrum* Form II Rubisco (PDB 5RUB) (*46*). In the expanded CAVER cohort, GEN_10’s static cavity descriptors remained within the natural representative range, and no descriptor–activity or descriptor–apparent-specificity relationship survived false-discovery-rate correction (*44*). The structure therefore rules out gross fold, assembly, or binding-pocket collapse as the explanation for low activity but does not identify a causal mechanism.

**Figure 4.**
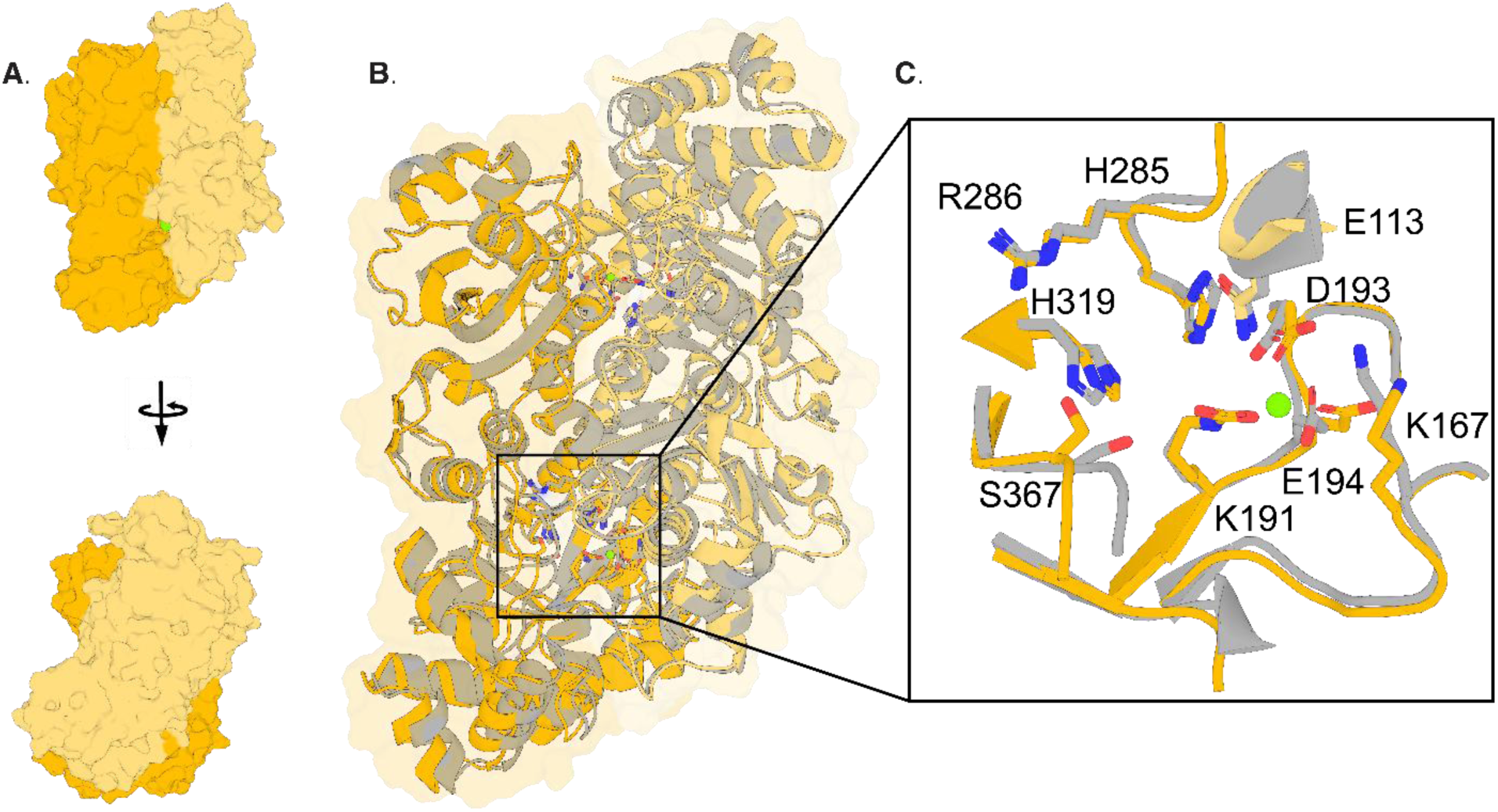
The GEN_10 crystal structure reproduces the predicted dimer and active-site geometry. **(A)** The 1.40 Å GEN_10 structure (gold) forms an L_2_ dimer. **(B)** Superposition with the AlphaFold model (gray) gives Cα RMSDs of 0.747 Å for the monomer and 0.621 Å for the dimer. Mg^+2^ is green. **(C)** Active-site superposition gives a Cα RMSD of 0.557 Å and shows the aligned catalytic pocket surrounding Mg^+2^.

## Discussion

RubisGen allows a broad search across Rubisco sequence space. Of the 21 representative sequences evaluated, five produced 3PGA at levels within the range of the natural enzymes used for training, and their apparent specificity estimates spanned a similarly broad range (**Figure 3C, D; Tables S3–S4**). The activity assay is what establishes that these designs are catalytically active; the crystal structure of GEN_10 provides independent evidence that a design from this new region of sequence space folds into the expected Rubisco dimer (**Figure 4**).

The selection funnel converted approximately 5.6 million generated sequences into 21 experimental candidates by combining complementary sequence and structural filters (**Figure 1; Table S1**). Because no original selection score subsequently predicted soluble recovery after multiple-testing correction (**Table S7**), its value was in organizing a broad, auditable search rather than acting as a single predictor of function. The designs that emerged were also substantially diverged from the natural collection they were compared against: the five active designs shared 70.6–80.4% identity with their nearest natural match, and their differences from that match were spread across the sequence rather than confined to a few long blocks (**Figures S1–S3; Tables S11a and S11b**).

A central realization from GEN_10 is that preservation of the Rubisco fold is separable from catalytic optimization. Its crystal structure recapitulated the predicted L_2_ assembly and active-site geometry despite 39.2% identity across the full gapped alignment, and 80.0% of the eligible rigid active-site positions retained amino acid identity (**Figure 4; Table S10**). Yet neither gross structural deviation, interface conservation, nor static cavity geometry explained the measured activity range (**Figure S5; Tables S7–S8**). This strengthens the view that Rubisco performance depends on context-dependent interactions beyond a minimal catalytic motif and is consistent with evidence that PLMs can preserve higher-order evolutionary constraints (*24*, *27*, *28*). The immediate path forward is to pair generative sampling with kinetic and perturbational measurements that can resolve activation, conformational dynamics, and epistasis.

Soluble recovery appears to impose a distinct upstream constraint. The only FDR-supported computational association was a lower signed basic-minus-acidic residue fraction in the six soluble designs, whereas global predicted structural similarity and the original selection metrics were not significant. This compositional proxy is consistent with an electrostatic contribution to heterologous recovery. It is not a formal-charge calculation, and with six soluble designs the present cohort establishes a direction rather than a threshold, which prospective designs can test directly while preserving active-site and interface constraints.

In conclusion, RubisGen enables generation and exploration of Rubisco sequence space at a scale previously inaccessible through directed evolution or database mining. The resulting sequences encoded soluble proteins with measurable melting temperatures, apparent 3PGA and 2PG activity within the assayed natural range, an apparent specificity estimate beyond the measured natural representative range, and sub angstrom agreement between a predicted and experimental dimer. These results support a discovery framework in which broader sequence generation, experimental feedback, and model refinement progressively enrich the rare combinations of activity, specificity, folding, and assembly required to improve biological carbon fixation.

## Methods

### Dataset curation and model training

ProGen-2 (*25*) was fine-tuned on a curated collection of sequences defined as bona fide Rubiscos (*1*). Forms I and RLPs and proteins outside 361–500 residues were excluded, leaving 3,028 sequences. Data were divided 80:10:10 into training, validation, and test sets. Training used an A100 GPU, a learning rate of 2 × 10^−4^, local and global batch sizes of 8 and 32, respectively, and a patience of 500 steps.

We trained a discriminator following a published generative–discriminative strategy (*27*). Its dataset contained 36,276 natural Rubiscos after excluding RLPs (*1*) and 36,998 generated sequences sampled at top-*p* = 1 and temperature = 1; all sequences were 200–500 residues. The generator set was more stringently curated because coherent full-length sequences were required for generation. The discriminator used a broader length-filtered set, including fragments and Form I sequences, because its embedding-based natural-versus-generated classification task did not require every sequence to satisfy the generator’s full-length or form-specific curation criteria. A 90:5:5 training, validation, and test split was used to fine-tune ESM-2 (esm2_t6_8M_UR50D) (*26*) with the generator hyperparameters.

### Sequence generation and filtering

Because top-*p* = 0.25 produced repetitive sequences, subsequent analyses used 3.3 million, 1.2 million, and 1.1 million sequences sampled at top-*p* = 0.5, 0.75, and 1.0, respectively. Candidates were scored by the generator and discriminator and aligned to the training sequences with BLASTP (*47*). The discriminator returned p(generated); sequences with p(generated) > 0.5 were removed. The remainder were binned by natural-sequence identity (0–30%, 30–40%, 40–50%, 50–60%, 60–70%, or 70–100%); within each bin, candidates with the lowest p(generated) and lowest generator loss were retained independently. Sequences containing more than 10 consecutive identical residues were removed, followed by MMseqs2 clustering at 80% identity (*32*) and alignment against UniRef90 (*48*).

Dimeric structures of the remaining 1,375 sequences were predicted with AlphaFold multimer (*33*) and searched against the PDB with Foldseek (*34*). The experimental set was assembled from candidates above 70% UniRef90 identity after requiring mean pLDDT greater than 90, excluding named Form I and RLPs Foldseek hits, ranking by low p(generated) or generator loss, and manually inspecting active-site completeness. Additional structural triage used p(generated) ≤ 0.05, generator loss ≤ 0.7, ProteinMPNN scores at or above the candidate median, normalized SAP ≤ 0.5, and manual evaluation of active-site residues, termini, and loops. The final 21 designs spanned the retained sequence and structure-score ranges rather than representing the top value of any single metric (Table S1).

### Sequence similarity network and Sequence Analysis

We followed the procedures of constructing SSNs as outlined in a previous publication(*49*). Briefly, the SSN construction involves three key steps: First, collecting input sequences; second, analyzing these sequences for phylogenetic information using EFI-EST; and third, visualizing the results(*49*, *50*). Sequences acquired from both Prywes et al. 2023 and the generation of sequences described above were used as the input for EFI-EST analysis of Rubisco sequence space. The workflow contained 1,375 generated sequences, which were reduced to 987 centroids at 70% sequence identity and combined with 639 centroids from the natural non-Rubisco-like-protein sequence set (1,626 sequences total). The EFI-EST full network contained 1,626 nodes at an alignment-score threshold of 70, and the 40%-identity representative-node network contained 816 nodes. Generated sequences were labeled separately, natural nodes were annotated by Rubisco clade, and the network was visualized in Cytoscape (*51*).

### Phylogenetic tree construction

For the displayed phylogeny, the combined 1,626-sequence set was reduced to 342 centroids at 40% sequence identity. MAFFT produced an amino-acid alignment using --auto and --reorder (*40*), and trimAl with -gt 0.1 (*52*). FastTree then inferred an approximately maximum-likelihood tree from the trimmed alignment; branch support values were retained from the FastTree output, and the tree was displayed unrooted (*53*).

### Expression and purification of recombinant proteins

Synthetic genes encoding the 21 generated designs were synthesized by Twist Bioscience, and plasmids encoding the natural representatives were taken from the previously described plasmid library(*5*). Plasmids encoding N-terminal His_14_-bdSUMO-tagged RbcL were co-transformed into chemically competent BL21(DE3) Star *E. coli* with pBADES/EL. Cells were grown at 37 °C to OD_600_ = 0.7–0.8; GroEL/ES expression was induced with 0.2% (w/v) arabinose for 2 hours. Cells were transferred to fresh terrific broth without arabinose, induced with 1 mM IPTG, and shaken for 16 hours at 16 °C. Pellets were resuspended in lysis buffer [20 mM sodium phosphate (pH 8.0), 300 mM NaCl, 10 mM imidazole, 5% glycerol, and 2 mM MgCl_2_] containing approximately 5 mM PMSF, subjected to one freeze–thaw cycle at −80 °C, and sonicated. Soluble material was collected at 15,000 × g for 20 min and applied to equilibrated Ni–NTA resin. After washing, resin was resuspended in bdSENP1 buffer [20 mM HEPES (pH 8.0), 300 mM NaCl, and 10% glycerol] and incubated overnight at 4 °C with bdSENP1 to remove the His_14_-bdSUMO tag. Purity was validated via SDS-PAGE.

### Protein thermal-shift assay

Samples contained 1 mg mL^−1^ protein, 1× Protein Thermal Shift phosphate buffer, and 4× dye (Thermo Fisher) in MicroAmp optical strips. A QuantStudio 3 instrument held samples at 16 °C for 1 min, increased the temperature to 100 °C at 0.05 °C s^−1^, and held at 100 °C for 1 min. Melting temperatures were calculated with Protein Thermal Shift Software.

### LC-MS-based activity assay

Rubisco variants were assayed in 200 µL reaction mixtures containing 50 mM HEPES (pH 8.0). Before initiation, enzyme-containing mixtures were incubated for 20 min at 30 °C under 5% CO_2_ and 21% O_2_ to activate Rubisco. Reactions were initiated by adding ribulose-1,5-bisphosphate (RuBP) to 1 mM and were incubated for 24 hours at 30 °C under the same atmosphere.

After incubation, 40 µL aliquots were mixed with 50 µL derivatization reagent containing 80 mM 3-nitrophenylhydrazine (3-NPH), 48 mM 1-ethyl-3-(3-dimethylaminopropyl)carbodiimide (EDC), and 2.4% (v/v) pyridine, adapted from Hofstetter et al. (2023)(*54*). Derivatization was simultaneously performed for 30 min at 18 °C to optimize 3PGA derivatization, and for 24 hours at 60 °C to optimize 2PG derivatization. Derivatized samples were filtered through Millipore MultiScreen Solvinert plates by centrifugation at 3,000 × g for 10 min at 4 °C.

Derivatized products were quantified by LC-MS, modified from Wu et al. (2026)(*55*). Samples (1 µL) were analyzed using an Agilent 1260 liquid chromatography system fitted with a Puranalyzed using18 endcapped column (50 mm × 2.1 mm, 2 µm particle size; Merck/Supelco). The mobile phases were 0.1% (v/v) formic acid in water (solvent A) and 0.1% (v/v) formic acid in acetonitrile (solvent B). Separation was performed at 0.3 mL min^−1^ using the following gradient: 0–2 min, 20% B; 2–5 min, 20–50% B; 5–8 min, 50–95% B; 8–10 min, 95% B; 10–13 min, 95–20% B; and 13–16 min, 20% B. The column temperature was maintained at 60 °C.

The LC system was coupled to an Agilent 6120 single-quadrupole mass spectrometer equipped with an ESI source and operated in negative-ion selected-ion-monitoring (SIM) mode. Source settings were: capillary voltage, 5,200 V; drying-gas flow rate, 13 L min^−1^; nebulizer pressure, 60 psig; and drying-gas temperature, 350 °C. Ions at m/z 425 and 455 were monitored for the derivatized 2PG and 3PGA products, respectively.

Analyte concentrations were determined from integrated LC–MS peak areas using calibration curves prepared from authentic 2PG and 3PGA standards that were derivatized identically to the samples. Calibration ranges were analyte-and plate-specific, and the lowest quantified 2PG standard used for each reporting threshold is listed in Table S5. Product concentrations were back-calculated to their concentrations in the original reaction mixtures. Product amounts and apparent product-formation activities were calculated independently for each analyte as:

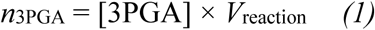

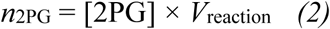

where n3PGA and n2PG denote the amount of 3PGA and of 2PG formed in the reaction, in micromoles; [3PGA] and [2PG] are the measured product concentrations in the original reaction mixture, in millimolar; and Vreaction is the reaction volume, 200 µL. Apparent product-formation activity (A3PGA; A2PG; µmol product mg protein^−1^ min^−1^) was calculated as:

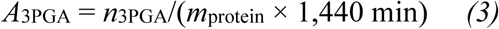

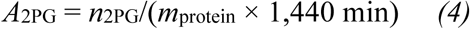

Here A3PGA and A2PG are the apparent product-formation activities, mprotein is the mass of purified protein in the reaction, in milligrams, and 1,440 min is the 24-hour reaction time over which product accumulated.

### Apparent 2PG activity reporting and specificity calculation

Apparent 2PG specific activity was reported only for enzymes with at least two technical replicates meeting the S_C/O,app_ reporting criterion. Apparent specificity is an endpoint quantity derived from the products that accumulated over the whole reaction; it is not the conventional kinetic specificity factor, which is defined from carboxylation and oxygenation rate constants, and the two should not be compared directly. For those enzymes, the enzyme-level mean and SD used all available original 2PG technical replicates, regardless of whether each replicate individually passed the specificity threshold. Entries without reportable enzyme-level apparent specificity were assigned

N.D. for apparent 2PG activity, indicating that raw 2PG did not exceed the applicable plate-specific reporting threshold defined below. Each technical replicate was matched to plate-level RuBP negative controls for the apparent specificity analysis. The matched control mean was subtracted from raw 2PG. 3PGA was not corrected for background because no background 3PGA was detected in the negative controls. The final raw-2PG threshold was the larger of the control mean + 2 SD and the lowest quantified 2PG standard, and raw sample 2PG had to be strictly greater than this value. Corrected 2PG concentration entered the apparent specificity calculation only when the raw value passed that reporting threshold; corrected 2PG concentration then had to be finite and positive, and 3PGA had to be finite and non-negative. The 0.5 factor accounts for two 3PGA molecules per carboxylation event and one 3PGA plus one 2PG per oxygenation event. The product ratio and the apparent specificity were calculated with Equations (5) and (6), in which vc and vo are the carboxylation and oxygenation velocities and 0.1976 is the dissolved-gas ratio at 30°C, 5% CO_2_, and atmospheric O_2_. Enzyme-level apparent specificity means and SDs were reported only when at least two technical replicates passed all criteria. N.C. denotes an apparent specificity estimate not calculated because the specificity-reporting criteria were not met; it does not denote zero specificity.

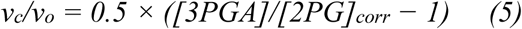

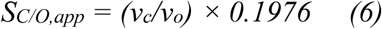

### Crystallization and structure determination

Ni-NTA–purified Rubisco was further subjected to size-exclusion chromatography (SEC) on a Superose 6 Increase 10/300 GL column in a final buffer containing 100 mM Hepes (pH 8), 100 mM NaCl, 25 mM MgCl2, 5 mM NaHCO3, and 1 mM DTT. GEN_10 was screened against the following crystallization screens: MCSG-1 (Anatrace); Crystal Screen, SaltRx, PEG/Ion, Index, and PEGRx (Hampton Research); and Berkeley Screen (*56*). GEN_10 crystals were obtained in 0.1 M HEPES pH 7.5 and 40% v/v Polyethylene glycol 400 and were cryoprotected with reservoir solution containing 20% (v/v) glycerol before flash-cooling in liquid nitrogen.

The X-ray dataset for GEN_10 was collected at the Berkeley Center for Structural Biology beamline 8.2.1 at the Advanced Light Source at Lawrence Berkeley National Laboratory. Diffraction data were processed using xia2 (*57*). The crystal structure of GEN_10 was solved by molecular replacement with the program PHASER (*58*) using the search model generated by AlphaFold (*33*). The atomic positions obtained from the molecular replacement were used to initiate model building using phenix.autobuild within the Phenix suite (*59*, *60*). Structure refinement was performed using the phenix.refine program (*61*). Manual rebuilding was done using COOT (*62*). Root-mean-square deviations from ideal geometries for bond lengths, angles, and dihedrals were calculated with Phenix (*60*). The stereochemical quality of the final model of GEN_10 was assessed with MolProbity (*63*). A summary of crystal parameters, data collection, and refinement statistics can be found in Table S9.

### Active-site sequence–structure mapping

To compare sequence conservation around the catalytic pocket, the activated *R. rubrum* Rubisco structure (PDB 9RUB) was used to define rigid reference residues within 8 Å of RuBP or Mg^+2^. Loop 6 and C-terminal lid residues were excluded because of ligand-state-dependent mobility. The 9RUB and GEN_10 sequences were aligned using an affine-gap BLOSUM62 alignment (*64*). Thirty-five reference residues met the rigid-anchor definition after the flexible positions were excluded; robust outlier rejection retained 31 mapped inliers for the GEN_10 local fit. A robust global Cα fit was followed by this local active-site fit; Mg^+2^, ligands, waters, and other heteroatoms in GEN_10 were excluded from fitting. Sequence identity was calculated as exact amino acid matches divided by the full gapped alignment length. Regional identities were calculated among aligned positions for operational 9RUB residue intervals 1–150, 151–420, and 421 through the C terminus. Carbamylated lysine was counted as lysine when evaluating residue identity. Both active sites were evaluated, but their sequence mappings are symmetry-equivalent technical checks rather than independent observations.

### Sequence–structure–function analysis

Forty proteins (21 generated and 19 natural representatives) were analyzed using the functional classifications and endpoint values reported in this manuscript. Six generated designs yielded soluble protein; five generated and 18 natural representative proteins yielded quantifiable 3PGA. Apparent 2PG activity and apparent specificity were available for 13 proteins. N.D. and N.C. entries were retained as missing and were never assigned zero. Quantitative relationships were stratified by generated versus natural representative proteins where sample size permitted.

Structural analyses used curated sequence-matched snapshots for all 21 designs and 17 natural representative proteins. *R. rubrum* and *Gallionella sp.* used experimental PDB 9RUB and 12TE respectively. Both 470-residue 4_10 chains were retained, with crystallographic KCX 200 represented as the underlying lysine in sequence comparisons. The experimental GEN_10 structure was held out for prediction-versus-experiment validation.

Sequences were aligned with MAFFT v7.505 using --auto(*40*). A phenotype-independent ligand-contact set was defined from activated *R. rubrum* Rubisco (PDB 9RUB) as protein residues having any heavy atom within 4.5 Å of RuBP, Mg^+2^, or formate. A strict screen required the same non-gap residue in all 23 3PGA-positive proteins followed by a different residue or gap in GEN_5 or *Ca. Woesearchaeota*. To reduce dependence on primary-sequence registration, each structure was also aligned to 9RUB using a coordinate-refined monotonic Cα mapping, and whole monomers and dimers were compared independently with US-align version 20260527 in sequence-independent mode (*42*). Dimer comparisons retained both subunits and optimized homodimer chain assignment.

Interchain residue contacts were defined by any non-hydrogen atom pair within 5.0 Å. Hydrophobic pairs used a 5.0 Å cutoff and oppositely charged residue pairs used a 4.0 Å cutoff. Active-site geometry was measured after a core Cα fit excluding ligand-contact positions.

Generated-design soluble recovery was compared across global sequence composition, predicted structural descriptors, original selection metrics, and whole-protein similarity to the natural panel. The signed basic-minus-acidic residue fraction was defined as (K+R+H−D−E)/sequence length and is a composition proxy rather than a formal charge or pI calculation. Two-sided Mann–Whitney tests, group medians, and Cliff’s delta were reported.

The curated-coordinate CAVER analysis used 18 structures matched to the primary cohort. Both active sites were mapped to a common 9RUB frame, analyzed separately, and averaged before protein-level testing. CAVER used a 0.9 Å probe and 4.0 Å shell, with a 3.0 Å shell sensitivity analysis (*44*). Standardized cavity electrostatics were calculated at pH 7 using PDB2PQR, PROPKA, and APBS (*65*, *66*). Primary Spearman tests used 100,000 deterministic permutations and 10,000 bootstrap resamples. Benjamini–Hochberg correction was applied within each predefined statistical family(*67*). Static cavities were interpreted as geometric descriptors, not evidence of substrate transport or protein dynamics.

Across all analyses, absence of an FDR-supported association was not interpreted as equivalence or proof that a feature is mechanistically irrelevant. No classifier, feature-selection model, or predictive threshold was fit because only six designs were soluble and five generated designs had quantifiable 3PGA.

### Sequence novelty analysis

Identity to the nearest natural sequence was taken, for each of the 1,375 structurally evaluated designs, from the original sequence comparison against the curated natural Rubisco collection used for training(*47*). Designs returning no match were reported as a separate category and were never assigned zero identity. These values are a comparison against that curated collection and are a different measurement from the UniRef90 identities reported in Tables S1 and S4.

For the two-dimensional sequence map, the same set of sequences used for the similarity network (987 generated and 639 natural representative sequences) was aligned with MAFFT(*40*), reduced to 50 principal components, and projected with UMAP using 30 neighbours, a minimum distance of 0.1, a Euclidean metric, and a fixed random seed(*68*). Stability was checked across three random seeds and neighbour settings of 15, 30, and 50. The map is a qualitative picture of how the sequences are arranged and was not used as a numerical measure of novelty.

To test whether the designs resemble mosaics of long natural segments, the 21 experimentally tested designs were searched against the same natural collection with MMseqs2, retaining up to 50 matches per design(*1*, *32*). Matches were grouped at 90% identity before the cumulative identical-residue coverage of the closest one, two, three, and five groups was calculated, together with the longest run of consecutive identical residues, the highest identity in any 30-residue window, and the longest merged region in which overlapping 30-residue windows were at least 90% identical. The clustering of identical residues within each design’s closest match was compared with 10,000 random rearrangements that preserved the observed numbers of matching and mismatching positions, and the resulting P values were corrected across the five active designs with the Holm procedure(*69*). No coverage filter was applied; for the five active designs the closest match covered 91.1–99.5% of the design and 91.7–97.9% of the natural sequence. Per-design values are given in Tables S11a and S11b.

## Acknowledgements

The authors wish to extend their sincere gratitude to their families for their steadfast support and encouragement provided throughout the course of this work. We are deeply indebted to our partners, parents, and loved ones, whose patience and understanding sustained us during the demands of this research. Their contributions, though not visible in the data presented herein, are inseparable from its completion.

## Contributions

A.J.K., S.K.S.C J.B.S., P.M.S. carried out overall experimental design. A.J.K and S.K.S.C carried out dataset creation, model training, generation and filtering of generated sequences. A.J.K, R.Z.W. and MG carried out LCMS assay development. A.J.K carried out all analysis. All protein samples were prepared by A.J.K. *E. coli* transformations, growth and expression were carried out by J.L. conducted all protein thermal shift experiments. J.H.P., P.D.A. performed x-ray crystallography data acquisition, image processing, and structure determination. All authors contributed to writing and manuscript preparation.

