## Supplemental_Information for "De novo Rubisco design with protein language models"

### Supplementary information:

#### Supplementary Figures:

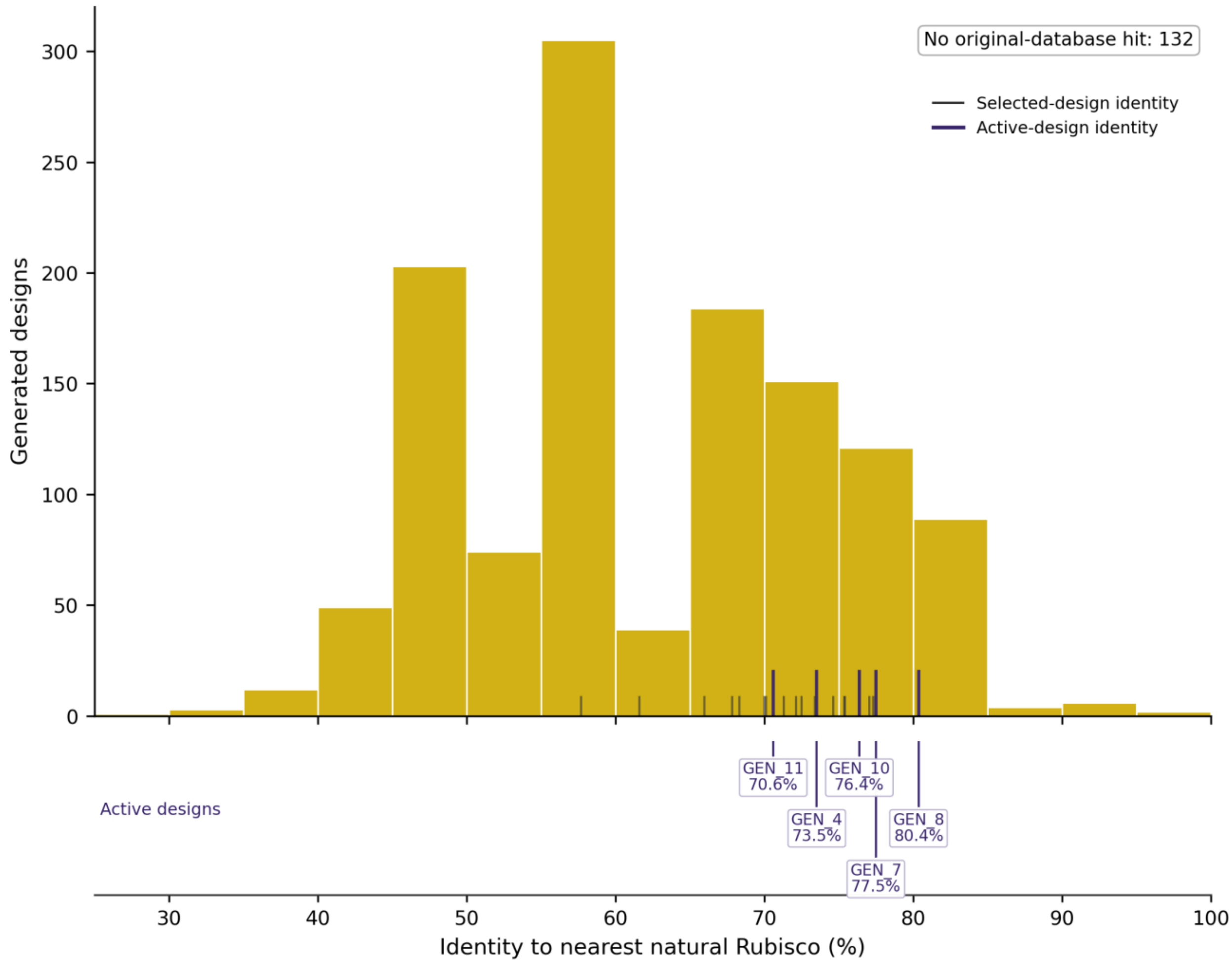


Figure S1. Identity of generated designs to their nearest natural Rubisco. Distribution of nearest-natural sequence identity for the 1,243 of 1,375 structurally evaluated designs that returned a match against the curated natural Rubisco collection. The median is 59.73% and the interquartile range is 54.02–72.09%. The 132 designs with no match are counted separately and are not shown as zero identity.

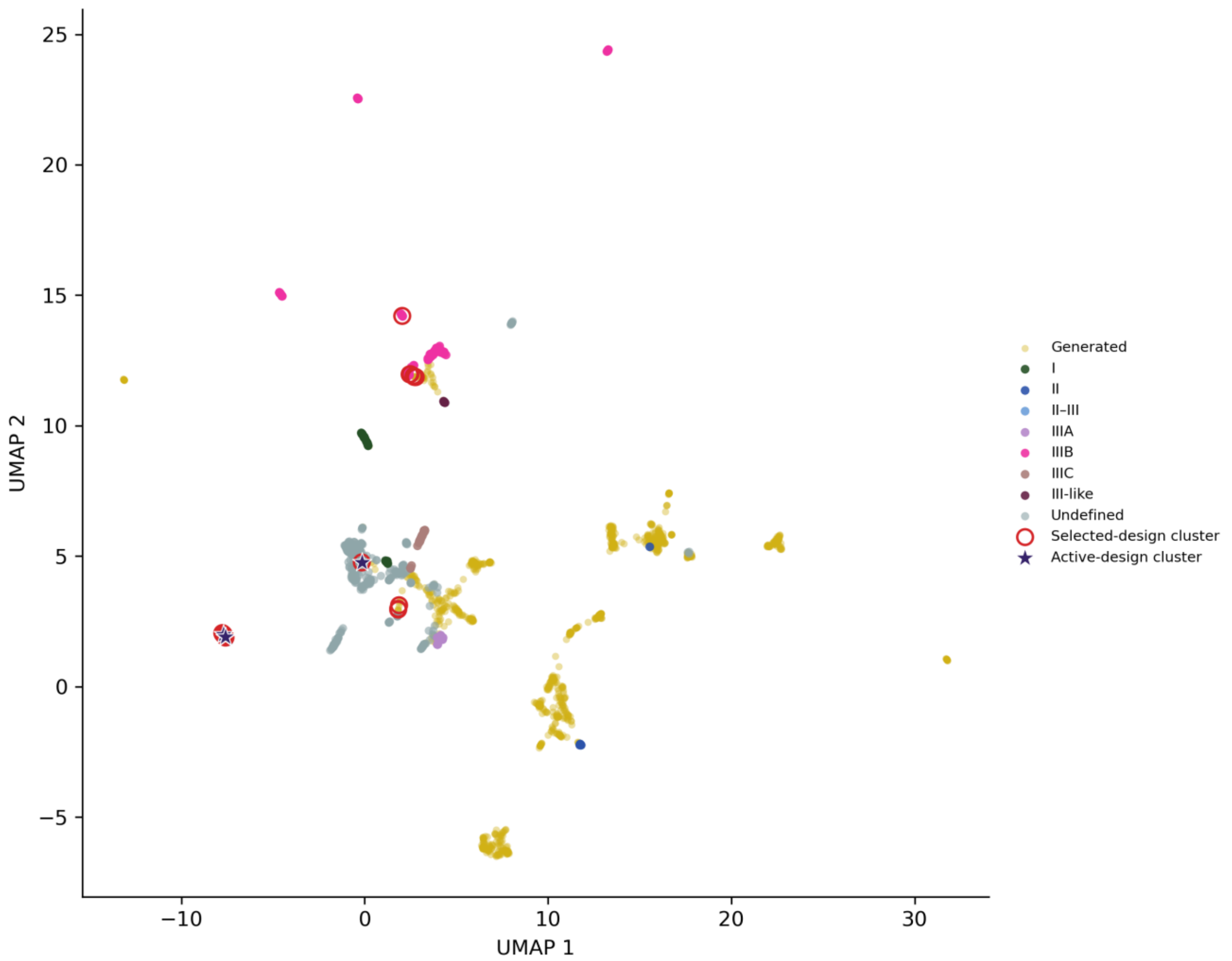


Figure S2. Two-dimensional sequence map of generated and natural Rubiscos. UMAP projection of the aligned sequence-similarity-network set (987 generated and 639 natural representative sequences). Natural sequences are coloured by Rubisco form using the palette of Figure 2A, with sequences that are unassigned in the source metadata shown as undefined. The projection is a qualitative visualization and is not used as a quantitative measure of novelty.

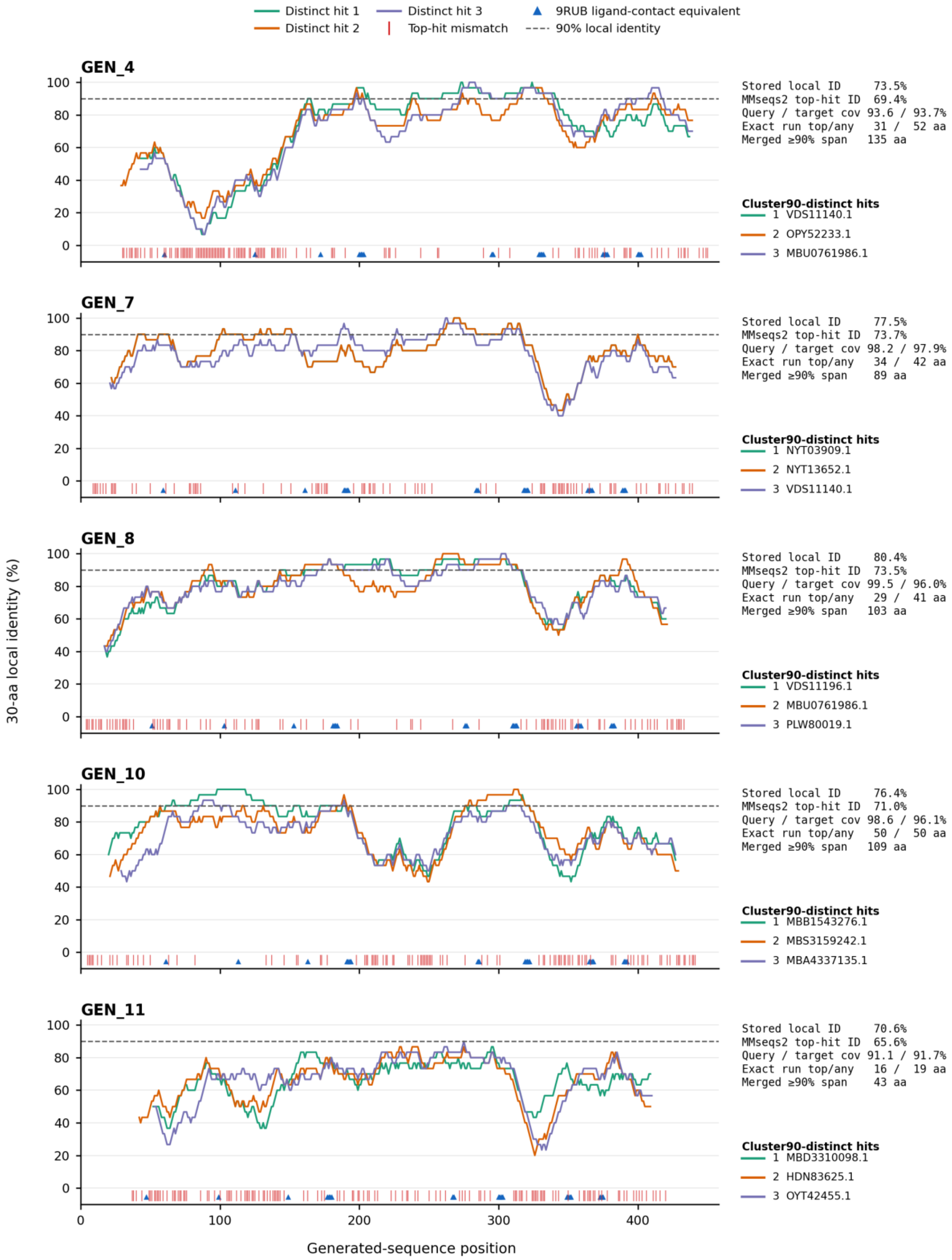

Distinct hit 1
Distinct hit 2
Distinct hit 3
Top-hit mismatch
9RUB ligand-contact equivalent
90% local identity
GEN_4
Stored local ID 73.5%
MMseqs2 top-hit ID 69.4%
Query / target cov 93.6 / 93.7%
Exact run top/any 31 / 52 aa
Merged ≥90% span 135 aa
Cluster90-distinct hits
1 VDS11140.1
2 OPY52233.1
3 MBU0761986.1
GEN_7
Stored local ID 77.5%
MMseqs2 top-hit ID 73.7%
Query / target cov 98.2 / 97.9%
Exact run top/any 34 / 42 aa
Merged ≥90% span 89 aa
Cluster90-distinct hits
1 NYT03909.1
2 NYT13652.1
3 VDS11140.1
GEN_8
Stored local ID 80.4%
MMseqs2 top-hit ID 73.5%
Query / target cov 99.5 / 96.0%
Exact run top/any 29 / 41 aa
Merged ≥90% span 103 aa
Cluster90-distinct hits
1 VDS11196.1
2 MBU0761986.1
3 PLW80019.1
GEN_10
Stored local ID 76.4%
MMseqs2 top-hit ID 71.0%
Query / target cov 98.6 / 96.1%
Exact run top/any 50 / 50 aa
Merged ≥90% span 109 aa
Cluster90-distinct hits
1 MBB1543276.1
2 MBS3159242.1
3 MBA4337135.1
GEN_11
Stored local ID 70.6%
MMseqs2 top-hit ID 65.6%
Query / target cov 91.1 / 91.7%
Exact run top/any 16 / 19 aa
Merged ≥90% span 43 aa
Cluster90-distinct hits
1 MBD3310098.1
2 HDN83625.1
3 OYT42455.1
30-aa local identity (%)
Generated-sequence position


18

Figure S3. Position-resolved comparison of the five active designs with their closest natural matches. For GEN_4, GEN_7, GEN_8, GEN_10, and GEN_11, tracks show, along the length of each design, the positions that are identical to the closest natural match, the local identity in a sliding 30-residue window, and the positions equivalent to ligand-contact residues. Numerical values for all 21 designs are given in Tables S11a and S11b.

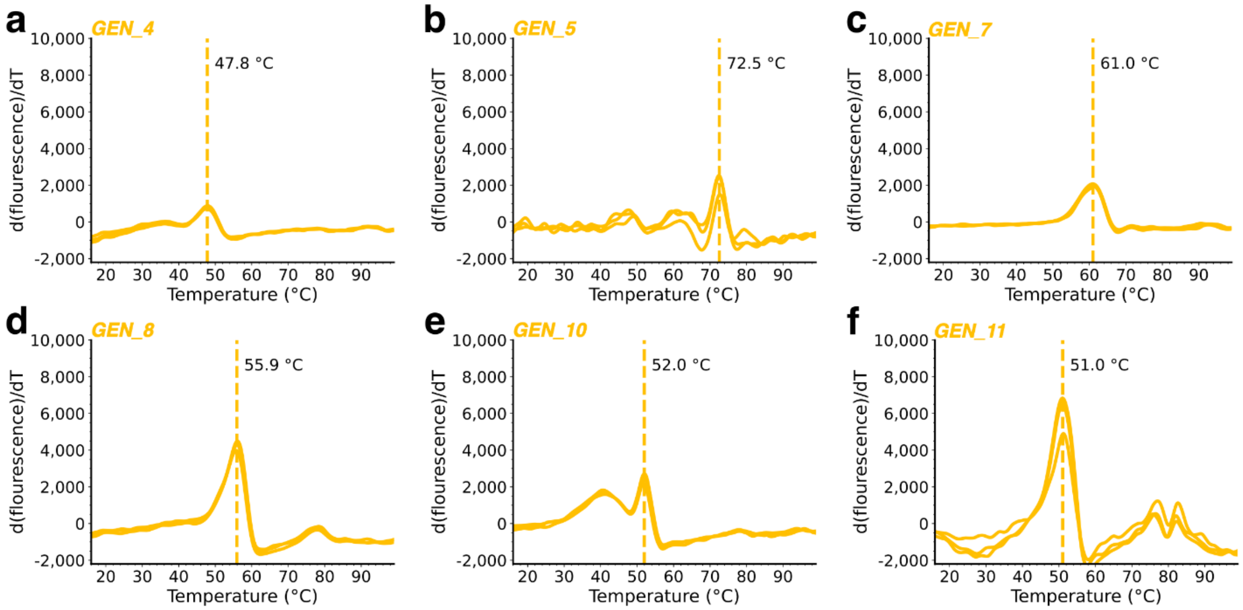


Figure S4. Melting profiles of soluble generated Rubiscos. Protein thermal-shift traces are shown with annotated mean melting temperatures ($T_m$) from three technical replicates. (A) GEN_4. (B) GEN_5. (C) GEN_7. (D) GEN_8. (E) GEN_10. (F) GEN_11.

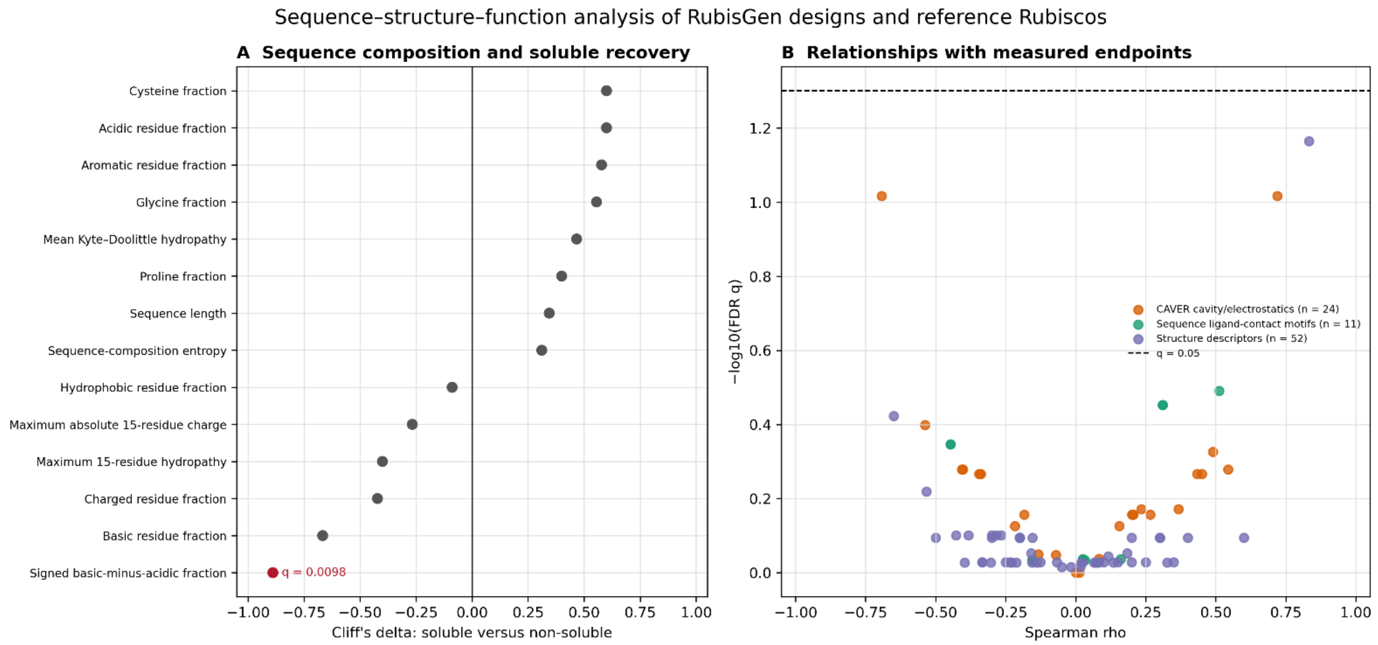

Figure S5. Sequence–structure–function analysis. (A) Cliff's delta for predefined global sequence-composition features comparing six soluble with 15 non-soluble RubisGen designs. Red denotes $q < 0.05$ after Benjamini–Hochberg correction within the predefined family. The signed basic-minus-acidic residue fraction was the only FDR-supported feature. (B) Spearman effect sizes and FDR values for predefined ligand-contact, static structure, and primary CAVER relationships with apparent 3PGA activity, apparent 2PG activity, apparent specificity, or melting temperature, as applicable. No relationship in panel B passed $q < 0.05$. Complete test definitions and sample sizes are provided in Tables S7 and S8.

**Supplementary tables:**

Table S1. Unified candidate-selection metrics and top Foldseek PDB hits for the 21 experimentally tested RubisGen designs. p(generated) is the ESM-2 discriminator probability assigned to the generated class; lower values are more natural-like. Foldseek identity is reported for the top PDB hit. Soluble recovery is the experimental outcome and was not a selection filter.

| Design | p(generated) | Generator loss | UniRef90 identity (%) | Mean pLDDT | Top PDB hit | Foldseek identity | Soluble |
|---|---|---|---|---|---|---|---|
| GEN_1 | 0.000795 | 0.771 | 77.3 | 92.80 | 5MAC | 0.660 | No |
| GEN_2 | 0.062246 | 0.365 | 80.2 | 94.50 | 6HUN | 0.474 | No |
| GEN_3 | 0.351393 | 0.391 | 82.7 | 95.19 | 6HUN | 0.460 | No |
| GEN_4 | 0.000877 | 0.392 | 73.5 | 90.33 | 5MAC | 0.593 | Yes |
| GEN_5 | 0.002686 | 0.398 | 79.6 | 93.70 | 5MAC | 0.621 | Yes |
| GEN_6 | 0.328086 | 0.402 | 82.2 | 92.60 | 6HUN | 0.457 | No |
| GEN_7 | 0.001253 | 0.445 | 77.5 | 94.60 | 5MAC | 0.690 | Yes |
| GEN_8 | 0.001242 | 0.461 | 80.4 | 94.82 | 5MAC | 0.694 | Yes |
| GEN_9 | 0.000959 | 0.852 | 75.4 | 90.94 | 5MAC | 0.610 | No |
| GEN_10 | 0.001203 | 0.473 | 76.4 | 94.28 | 5MAC | 0.665 | Yes |
| GEN_11 | 0.034869 | 0.609 | 70.6 | 90.55 | 6HUN | 0.515 | Yes |
| GEN_12 | 0.020308 | 0.561 | 74.6 | 93.10 | 3WQP | 0.564 | No |
| GEN_13 | 0.020860 | 0.665 | 72.5 | 90.84 | 5MAC | 0.614 | No |
| GEN_14 | 0.023596 | 0.558 | 79.6 | 92.70 | 6HUN | 0.423 | No |
| GEN_15 | 0.017305 | 0.509 | 75.4 | 92.37 | 8DHT | 0.603 | No |
| GEN_16 | 0.007944 | 0.583 | 70.1 | 94.52 | 2D69 | 0.558 | No |
| GEN_17 | 0.031439 | 0.551 | 74.9 | 94.86 | 2D69 | 0.543 | No |
| GEN_18 | 0.010288 | 0.542 | 73.0 | 93.69 | 5MAC | 0.590 | No |
| GEN_19 | 0.024956 | 0.516 | 72.5 | 92.61 | 3WQP | 0.498 | No |
| GEN_20 | 0.047037 | 0.546 | 70.0 | 94.32 | 3WQP | 0.488 | No |
| GEN_21 | 0.048498 | 0.697 | 73.4 | 94.55 | 2D69 | 0.518 | No |

Table S2. Enzyme metadata for the endpoint activity analysis. Sequence accessions or source identifiers are provided for the natural representative proteins; *Gallionella sp.* corresponds to the fast natural Form II $L_2$ Rubisco OGS68397.1, and *R. rubrum* is identified by UniProt P04718. Oligomeric-state assignments are carried over from published SEC–SAXS measurements; detailed SEC–SAXS tables are not repeated here. An *em* dash indicates that an assignment was unavailable or not applicable.

| Display name | Organism/source | Sequence accession/identifier | Rubisco form | Oligomeric state |
|---|---|---|---|---|
| GEN_10 | Generated Rubisco design | — | not applicable | L2 |
| GEN_11 | Generated Rubisco design | — | not applicable | — |
| GEN_4 | Generated Rubisco design | — | not applicable | — |

| | | | | |
|---|---|---|---|---|
| GEN_5 | Generated Rubisco design | — | not applicable | — |
| GEN_7 | Generated Rubisco design | — | not applicable | — |
| GEN_8 | Generated Rubisco design | — | not applicable | — |
| A. boonei | Aciduliprofundum boonei | WP_008086508.1 | unclassified | L2 |
| A. degensii | Ammonifex degensii KC4 | ACX52946.1 | Iβ | L2 |
| Altiarchaeales archaeon | Altiarchaeales archaeon | HIE34177.1 | unclassified | L2 |
| Bacterium NOY14888 | unclassified bacterium (NOY14888.1) | NOY14888.1 | unclassified | L2 |
| Bacterium RKZ25299 | unclassified bacterium (RKZ25299.1) | RKZ25299.1 | unclassified I | L2 |
| Candidatus Dojkabacteria | Candidatus Dojkabacteria | KKP92911.1 | II-III | L2 |
| *Ca. Woesearchaeota* | Candidatus Woesearchaeota | PIU30467.1 | unclassified | L2 |
| Candidatus Gottesmaniibacteriota | Candidatus Gottesmaniibacteriota | OGG30932.1 | II-III | L2 |
| F. fontis | Fervidicoccus fontis | PMB76193.1 | Iβ | L2 |
| Kerfeldbacteria bacterium | Kerfeldbacteria bacterium | MBI5465798.1 | III-like | L2 |
| Lokiarchaeota archaeon | Lokiarchaeota archaeon | Candidatus_Lokiarchaeota_archaeon_strain_HM5_B55 | unclassified | L2 |
| Methanobacteriota archaeon | Methanobacteriota archaeon | HDR53286.1 | unclassified | L2 |
| Patescibacteria sp. (MBU2523915.1) | Patescibacteria sp. (MBU2523915.1) | MBU2523915.1 | unclassified | L2 |
| Patescibacteria sp. (MBU1445715.1) | Patescibacteria sp. (MBU1445715.1) | MBU1445715.1 | unclassified | L2 |
| Prometheoarchaeum syntrophicum | Prometheoarchaeum syntrophicum | WP_147662334.1 | unclassified | L2 |
| R. faecalis | Rhodopseudomonas faecalis | WP_110780429.1 | II | L6 |
| Thermodesulfobium acidiphilum | Thermodesulfobium acidiphilum | WP_108308545.1 | Iβ | L2 |
| Gallionella sp. | Gallionella sp. | OGS68397.1 | II | L2 |
| R. rubrum | Rhodospirillum rubrum | P04718 | II | L2 |

Table S3. Natural representative Rubisco endpoint 3PGA activity, reportable apparent 2PG activity, and apparent specificity. Reactions were performed for 24 hours at 30 °C under 5% $CO_2$ and atmospheric $O_2$ in 200 µL with 1 mM RuBP. Apparent 3PGA specific activities are mean ± SD from three technical replicates on the selected source plate, except for Patescibacteria sp. (MBU2523915.1), for which one replicate was not quantifiable and the mean ± SD summarizes two technical replicates. Apparent 2PG activities are mean ± SD from all available original technical replicates and are reported only for enzymes with at least two technical replicates meeting the apparent specificity criterion; a two-replicate 2PG summary is marked. Apparent specificity values are mean ± SD from threshold-passing technical replicates; estimates based on two passing replicates are marked. N.D., 3PGA was not quantifiable or 2PG was not reliably detected under the applicable plate-specific reporting criterion, as applicable; N.C., apparent specificity was not calculated because the specificity-reporting criteria were not met.

| **Enzyme** | **Form** | **Apparent 3PGA specific activity (µmol mg⁻¹ min⁻¹)** | **Apparent 2PG specific activity (µmol mg⁻¹ min⁻¹)** | **Apparent specificity** | **Tm (°C)** |
|---|---|---|---|---|---|
| Gallionella sp. | II | $(1.84 \pm 0.05) \times 10^{-3}$ | $(8.74 \pm 0.74) \times 10^{-5}$ | 2.94 ± 0.42 | — |
| Bacterium RKZ25299 | unclassified I | $(8.89 \pm 0.19) \times 10^{-4}$ | $(1.07 \pm 0.36) \times 10^{-4}$ (n = 2) | 1.08 ± 0.54 (n = 2) | — |
| R. faecalis | II | $(4.59 \pm 0.08) \times 10^{-4}$ | $(8.06 \pm 1.00) \times 10^{-5}$ | 0.57 ± 0.10 | — |
| F. fontis | Iβ | $(3.82 \pm 0.23) \times 10^{-4}$ | $(3.18 \pm 0.08) \times 10^{-5}$ | 1.99 ± 0.03 | 78.3 ± 0.2 |
| Patescibacteria sp. (MBU1445715.1) | unclassified | $(3.41 \pm 0.37) \times 10^{-4}$ | $(3.59 \pm 0.09) \times 10^{-5}$ | 1.17 ± 0.17 | 63.7± 0.0 |
| R. rubrum | II | $(3.37 \pm 0.16) \times 10^{-4}$ | $(3.83 \pm 0.12) \times 10^{-5}$ | 1.13 ± 0.05 | 71.8 ± 0.1 |
| Kerfeldbacteria bacterium | III-like | $(2.82 \pm 0.15) \times 10^{-4}$ | $(2.29 \pm 0.12) \times 10^{-5}$ | 1.62 ± 0.21 | 68.7± 0.0 |
| Prometheoarchaeum syntrophicum | unclassified | $(2.43 \pm 0.16) \times 10^{-4}$ | $(5.26 \pm 0.56) \times 10^{-5}$ | 0.49 ± 0.09 | 61.6 ± 0.4 |
| Thermodesulfobium acidiphilum | Iβ | $(1.74 \pm 0.42) \times 10^{-4}$ | N.D. | N.C. | 77.7 ± 0.4 |
| Candidatus Dojkabacteria | II-III | $(1.07 \pm 0.36) \times 10^{-4}$ | N.D. | N.C. | — |
| A. degensii | Iβ | $(6.89 \pm 0.13) \times 10^{-5}$ | N.D. | N.C. | 95.2 ± 0.2 |
| Lokiarchaeota archaeon | unclassified | $(6.53 \pm 1.70) \times 10^{-5}$ | $(1.36 \pm 0.37) \times 10^{-5}$ | 0.96 ± 0.26 | 59.7± 0.3 |
| A. boonei | unclassified | $(5.98 \pm 0.28) \times 10^{-5}$ | $(1.33 \pm 0.02) \times 10^{-5}$ | 0.53 ± 0.01 | — |
| Bacterium NOY14888 | unclassified | $(5.12 \pm 0.18) \times 10^{-5}$ | $(1.51 \pm 0.10) \times 10^{-5}$ | 0.97 ± 0.17 (n = 2) | — |
| Altiarchaeales archaeon | unclassified | $(5.01 \pm 0.19) \times 10^{-5}$ | N.D. | N.C. | — |
| Patescibacteria sp. (MBU2523915.1) | unclassified | $(4.96 \pm 0.18) \times 10^{-5}$ | N.D. | N.C. | 57.5 ± 0.2 |
| Methanobacteriota archaeon | unclassified | $(3.46 \pm 0.05) \times 10^{-5}$ | N.D. | N.C. | 78.8± 0.2 |
| Candidatus Gottesmaniibacteriota | II-III | $(8.15 \pm 0.83) \times 10^{-6}$ | N.D. | N.C. | — |
| *Ca. Woesearchaeota* | unclassified | N.D. | N.D. | N.C. | — |

Table S4. Generated Rubisco endpoint 3PGA activity, reportable apparent 2PG activity, and apparent specificity. Reaction conditions are as in Table S3. Apparent 3PGA activities are mean ± SD from three technical replicates. Apparent 2PG activities are mean ± SD from all available original technical replicates and are reported only for designs with at least two technical replicates meeting the apparent specificity criterion. Relative 3PGA activity is calculated against the *R. rubrum* benchmark [(3.37 ± 0.16) × $10^{-4}$ µmol $mg^{-1}$ $min^{-1}$]. Apparent specificity values are mean ± SD from threshold-passing technical replicates; estimates based on two passing replicates are marked. N.D., 3PGA was not quantifiable or 2PG was not reliably detected under the applicable plate-specific reporting criterion, as applicable; N.C., apparent specificity was not calculated because the specificity-reporting criteria were not met.

| Design | % identity to nr90 | Apparent 3PGA specific activity (µmol $mg^{-1}$ $min^{-1}$) | Apparent 2PG specific activity (µmol $mg^{-1}$ $min^{-1}$) | Relative activity (%) | Apparent specificity | $T_m$ (°C) |
|---|---|---|---|---|---|---|
| GEN_11 | 70.60 | (3.08 ± 0.10) × $10^{-4}$ | (2.66 ± 0.20) × $10^{-5}$ | 91.4 | 1.51 ± 0.21 | 51.0 ± 0.15 |
| GEN_8 | 80.36 | (3.02 ± 0.13) × $10^{-4}$ | (1.69 ± 0.09) × $10^{-5}$ | 89.6 | 5.62 ± 0.37 (n = 2) | 55.9 ± 0.2 |
| GEN_7 | 77.50 | (2.25 ± 0.03) × $10^{-4}$ | N.D. | 66.8 | N.C. | 60.0 ± 0.3 |
| GEN_4 | 73.53 | (8.4 ± 3.4) × $10^{-6}$ | N.D. | 2.5 | N.C. | 47.8 ± 0.2 |
| GEN_10 | 76.39 | (4.37 ± 0.20) × $10^{-6}$ | N.D. | 1.3 | N.C. | 52.0 ± 0.4 |
| GEN_5 | 79.59 | N.D. | N.D. | N.D. | N.C. | 72.5 ± 0.2 |

Table S5. Plate-specific negative-control thresholds for the apparent specificity analysis. The final threshold is the larger of the control mean + 2 SD and the lowest quantified 2PG standard.

| Plate | Control 2PG mean (mM) | SD (mM) | n controls | Lowest standard (mM) | Final threshold (mM) |
|---|---|---|---|---|---|
| 042026 | 0.000000 | N.A. | 0 | N.A. | N.A. |
| 050526 | 0.053745 | 0.009671 | 6 | 0.030000 | 0.073087 |
| 051126 | 0.038661 | 0.013496 | 6 | 0.020000 | 0.065653 |
| 052026 | 0.049262 | 0.011666 | 6 | 0.020000 | 0.072594 |

Table S6. Technical-replicate 3PGA and 2PG quantification, apparent product-formation activities, and apparent specificity audit. Raw and corrected 2PG concentrations and threshold outcomes are retained for auditability. Apparent 2PG specific activity is reported only for enzymes with at least two valid apparent specificity replicates; all available original 2PG technical replicates are shown for eligible enzymes. N.Q., not quantifiable; N.D., 2PG was not reliably detected under the applicable plate-specific reporting criterion; N.C., apparent specificity was not calculated; N.A., not available.

| Display name | Replicate | Source plate | Protein (mg/mL) | 3PGA (mM) | Apparent 3PGA specific activity (µmol $mg^{-1}$ $min^{-1}$) | Raw 2PG (mM) | 2PG threshold (mM) | Corrected 2PG (mM) | Apparent 2PG specific activity (µmol $mg^{-1}$ $min^{-1}$) | Passed raw-2PG threshold | Apparent specificity |
|---|---|---|---|---|---|---|---|---|---|---|---|
| GEN_8 | 1 | 050526 | 3.08 | 1.285244 | 0.0002898 | 0.070327 | 0.073087 | 0.016581 | 1.5856 × $10^{-5}$ | No | N.C. |
| GEN_8 | 2 | 050526 | 3.08 | 1.330305 | 0.0002999 | 0.077815 | 0.073087 | 0.024069 | 1.7545 × $10^{-5}$ | Yes | 5.363 |
| GEN_8 | 3 | 050526 | 3.08 | 1.400695 | 0.0003158 | 0.076871 | 0.073087 | 0.023126 | 1.7332 × $10^{-5}$ | Yes | 5.887 |
| Gallionella sp. | 1 | 051126 | 1 | 2.571828 | 0.001786 | 0.133069 | 0.065653 | 0.094408 | 9.2409 × $10^{-5}$ | Yes | 2.593 |

| Display name | Replicate | Source plate | Protein (mg/mL) | 3PGA (mM) | Apparent 3PGA specific activity (µmol $mg^{-1}$ $min^{-1}$) | Raw 2PG (mM) | 2PG threshold (mM) | Corrected 2PG (mM) | Apparent 2PG specific activity (µmol $mg^{-1}$ $min^{-1}$) | Passed raw-2PG threshold | Apparent specificity |
|---|---|---|---|---|---|---|---|---|---|---|---|
| Gallionella sp. | 2 | 051126 | 1 | 2.716018 | 0.001886 | 0.130992 | 0.065653 | 0.092331 | $9.0967 \times 10^{-5}$ | Yes | 2.808 |
| Gallionella sp. | 3 | 051126 | 1 | 2.658935 | 0.001846 | 0.113576 | 0.065653 | 0.074916 | $7.8872 \times 10^{-5}$ | Yes | 3.409 |
| F. fontis | 1 | 051126 | 1.95 | 1.142795 | 0.000407 | 0.091970 | 0.065653 | 0.053310 | $3.2753 \times 10^{-5}$ | Yes | 2.020 |
| F. fontis | 2 | 051126 | 1.95 | 1.012955 | 0.0003607 | 0.087421 | 0.065653 | 0.048760 | $3.1133 \times 10^{-5}$ | Yes | 1.954 |
| F. fontis | 3 | 051126 | 1.95 | 1.060605 | 0.0003777 | 0.088845 | 0.065653 | 0.050184 | $3.1640 \times 10^{-5}$ | Yes | 1.990 |
| Kerfeldbacteria bacterium | 1 | 051126 | 4.08 | 1.608119 | 0.0002737 | 0.141814 | 0.065653 | 0.103154 | $2.4138 \times 10^{-5}$ | Yes | 1.442 |
| Kerfeldbacteria bacterium | 2 | 051126 | 4.08 | 1.598584 | 0.0002721 | 0.133799 | 0.065653 | 0.095139 | $2.2774 \times 10^{-5}$ | Yes | 1.562 |
| Kerfeldbacteria bacterium | 3 | 051126 | 4.08 | 1.755766 | 0.0002988 | 0.127794 | 0.065653 | 0.089133 | $2.1751 \times 10^{-5}$ | Yes | 1.848 |
| GEN_11 | 1 | 051126 | 3.548 | 1.511944 | 0.0002959 | 0.146359 | 0.065653 | 0.107699 | $2.8647 \times 10^{-5}$ | Yes | 1.289 |
| GEN_11 | 2 | 051126 | 3.548 | 1.595433 | 0.0003123 | 0.135119 | 0.065653 | 0.096459 | $2.6447 \times 10^{-5}$ | Yes | 1.536 |
| GEN_11 | 3 | 051126 | 3.548 | 1.611560 | 0.0003154 | 0.126441 | 0.065653 | 0.087780 | $2.4748 \times 10^{-5}$ | Yes | 1.715 |
| Patescibacteria sp. (MBU1445715.1) | 1 | 051126 | 2.884 | 1.241847 | 0.000299 | 0.153117 | 0.065653 | 0.114456 | $3.6869 \times 10^{-5}$ | Yes | 0.973 |
| Patescibacteria sp. (MBU1445715.1) | 2 | 051126 | 2.884 | 1.513980 | 0.0003646 | 0.145888 | 0.065653 | 0.107228 | $3.5129 \times 10^{-5}$ | Yes | 1.296 |
| Patescibacteria sp. (MBU1445715.1) | 3 | 051126 | 2.884 | 1.497902 | 0.0003607 | 0.148325 | 0.065653 | 0.109664 | $3.5715 \times 10^{-5}$ | Yes | 1.251 |
| R. rubrum | 1 | 050526 | 3.3 | 1.538064 | 0.0003237 | 0.175516 | 0.073087 | 0.121770 | $3.6935 \times 10^{-5}$ | Yes | 1.149 |
| R. rubrum | 2 | 050526 | 3.3 | 1.574560 | 0.0003313 | 0.185342 | 0.073087 | 0.131596 | $3.9003 \times 10^{-5}$ | Yes | 1.084 |
| R. rubrum | 3 | 050526 | 3.3 | 1.685511 | 0.0003547 | 0.185000 | 0.073087 | 0.131255 | $3.8931 \times 10^{-5}$ | Yes | 1.170 |
| Bacterium RKZ25299 | 1 | 050526 | 1.512 | 1.954913 | 0.0008979 | N.Q. | 0.073087 | N.C. | N.D. | No | N.C. |
| Bacterium RKZ25299 | 2 | 050526 | 1.512 | 1.887280 | 0.0008668 | 0.287837 | 0.073087 | 0.234091 | $1.3220 \times 10^{-4}$ | Yes | 0.698 |
| Bacterium RKZ25299 | 3 | 050526 | 1.512 | 1.962101 | 0.0009012 | 0.178265 | 0.073087 | 0.124520 | $8.1875 \times 10^{-5}$ | Yes | 1.458 |
| Bacterium NOY14888 | 1 | 050526 | 3.5 | 0.267732 | $5.312 \times 10^{-5}$ | 0.081616 | 0.073087 | 0.027870 | $1.6194 \times 10^{-5}$ | Yes | 0.851 |
| Bacterium NOY14888 | 2 | 050526 | 3.5 | 0.250319 | $4.967 \times 10^{-5}$ | 0.071156 | 0.073087 | 0.017411 | $1.4118 \times 10^{-5}$ | No | N.C. |
| Bacterium NOY14888 | 3 | 050526 | 3.5 | 0.255621 | $5.072 \times 10^{-5}$ | 0.075015 | 0.073087 | 0.021270 | $1.4884 \times 10^{-5}$ | Yes | 1.089 |
| Lokiarchaeota archaeon | 1 | 052026 | 4.972 | 0.398933 | $5.572 \times 10^{-5}$ | 0.081903 | 0.072594 | 0.032641 | $1.1440 \times 10^{-5}$ | Yes | 1.109 |
| Lokiarchaeota archaeon | 2 | 052026 | 4.972 | 0.394239 | $5.506 \times 10^{-5}$ | 0.081446 | 0.072594 | 0.032183 | $1.1376 \times 10^{-5}$ | Yes | 1.112 |
| Lokiarchaeota archaeon | 3 | 052026 | 4.972 | 0.609460 | $8.512 \times 10^{-5}$ | 0.127933 | 0.072594 | 0.078670 | $1.7868 \times 10^{-5}$ | Yes | 0.667 |
| R. faecalis | 1 | 051126 | 2.282 | 1.505723 | 0.0004582 | 0.302199 | 0.065653 | 0.263539 | $9.1964 \times 10^{-5}$ | Yes | 0.466 |
| R. faecalis | 2 | 051126 | 2.282 | 1.534172 | 0.0004669 | 0.239748 | 0.065653 | 0.201087 | $7.2959 \times 10^{-5}$ | Yes | 0.655 |

| Display name | Replicate | Source plate | Protein (mg/mL) | 3PGA (mM) | Apparent 3PGA specific activity (μmol $mg^{-1}$ $min^{-1}$) | Raw 2PG (mM) | 2PG threshold (mM) | Corrected 2PG (mM) | Apparent 2PG specific activity (μmol $mg^{-1}$ $min^{-1}$) | Passed raw-2PG threshold | Apparent specificity |
|---|---|---|---|---|---|---|---|---|---|---|---|
| R. faecalis | 3 | 051126 | 2.282 | 1.484318 | 0.0004517 | 0.252471 | 0.065653 | 0.213811 | $7.6831 \times 10^{-5}$ | Yes | 0.587 |
| A. boonei | 1 | 051126 | 6.98 | 0.591623 | $5.886 \times 10^{-5}$ | 0.132975 | 0.065653 | 0.094314 | $1.3230 \times 10^{-5}$ | Yes | 0.521 |
| A. boonei | 2 | 051126 | 6.98 | 0.578285 | $5.753 \times 10^{-5}$ | 0.131578 | 0.065653 | 0.092917 | $1.3091 \times 10^{-5}$ | Yes | 0.516 |
| A. boonei | 3 | 051126 | 6.98 | 0.631927 | $6.287 \times 10^{-5}$ | 0.136034 | 0.065653 | 0.097373 | $1.3534 \times 10^{-5}$ | Yes | 0.543 |
| Prometheoarchaeum syntrophicum | 1 | 052026 | 2.968 | 1.052309 | 0.0002462 | 0.198147 | 0.072594 | 0.148885 | $4.6362 \times 10^{-5}$ | Yes | 0.600 |
| Prometheoarchaeum syntrophicum | 2 | 052026 | 2.968 | 0.964658 | 0.0002257 | 0.231619 | 0.072594 | 0.182357 | $5.4194 \times 10^{-5}$ | Yes | 0.424 |
| Prometheoarchaeum syntrophicum | 3 | 052026 | 2.968 | 1.103807 | 0.0002583 | 0.244823 | 0.072594 | 0.195561 | $5.7283 \times 10^{-5}$ | Yes | 0.459 |
| GEN_7 | 1 | 050526 | 1.102 | 0.352593 | 0.0002222 | 0.064006 | 0.073087 | 0.010260 | N.D. | No | N.C. |
| GEN_7 | 2 | 050526 | 1.102 | 0.361327 | 0.0002277 | 0.061366 | 0.073087 | 0.007621 | N.D. | No | N.C. |
| GEN_7 | 3 | 050526 | 1.102 | 0.357705 | 0.0002254 | 0.055562 | 0.073087 | 0.001817 | N.D. | No | N.C. |
| Thermodesulfobium acidiphilum | 1 | 052026 | 1.958 | 0.398893 | 0.0001415 | 0.041542 | 0.072594 | -0.007720 | N.D. | No | N.C. |
| Thermodesulfobium acidiphilum | 2 | 052026 | 1.958 | 0.445906 | 0.0001581 | 0.045563 | 0.072594 | -0.003699 | N.D. | No | N.C. |
| Thermodesulfobium acidiphilum | 3 | 052026 | 1.958 | 0.625545 | 0.0002219 | 0.058014 | 0.072594 | 0.008751 | N.D. | No | N.C. |
| Candidatus Dojkabacteria | 1 | 052026 | 3.712 | 0.427072 | $7.99 \times 10^{-5}$ | 0.060890 | 0.072594 | 0.011627 | N.D. | No | N.C. |
| Candidatus Dojkabacteria | 2 | 052026 | 3.712 | 0.498350 | $9.323 \times 10^{-5}$ | 0.070234 | 0.072594 | 0.020972 | N.D. | No | N.C. |
| Candidatus Dojkabacteria | 3 | 052026 | 3.712 | 0.792724 | 0.0001483 | 0.099273 | 0.072594 | 0.050010 | N.D. | Yes | 1.468 |
| A. degensii | 1 | 050526 | 4.3 | 0.428270 | $6.916 \times 10^{-5}$ | 0.055393 | 0.073087 | 0.001648 | N.D. | No | N.C. |
| A. degensii | 2 | 050526 | 4.3 | 0.417620 | $6.745 \times 10^{-5}$ | 0.052727 | 0.073087 | -0.001018 | N.D. | No | N.C. |
| A. degensii | 3 | 050526 | 4.3 | 0.433657 | $7.004 \times 10^{-5}$ | 0.052400 | 0.073087 | -0.001345 | N.D. | No | N.C. |
| Altiarchaeales archaeon | 1 | 051126 | 3.828 | 0.285637 | $5.182 \times 10^{-5}$ | 0.042631 | 0.065653 | 0.003971 | N.D. | No | N.C. |
| Altiarchaeales archaeon | 2 | 051126 | 3.828 | 0.265239 | $4.812 \times 10^{-5}$ | 0.041451 | 0.065653 | 0.002790 | N.D. | No | N.C. |
| Altiarchaeales archaeon | 3 | 051126 | 3.828 | 0.277684 | $5.038 \times 10^{-5}$ | 0.043002 | 0.065653 | 0.004341 | N.D. | No | N.C. |
| Patescibacteria sp. (MBU2523915.1) | 1 | 050526 | 1.836 | N.Q. | N.Q. | 0.052666 | 0.073087 | -0.001079 | N.D. | No | N.C. |
| Patescibacteria sp. (MBU2523915.1) | 2 | 050526 | 1.836 | 0.127820 | $4.835 \times 10^{-5}$ | 0.051651 | 0.073087 | -0.002094 | N.D. | No | N.C. |
| Patescibacteria sp. (MBU2523915.1) | 3 | 050526 | 1.836 | 0.134366 | $5.082 \times 10^{-5}$ | 0.054396 | 0.073087 | 0.000650 | N.D. | No | N.C. |
| Methanobacteriota archaeon | 1 | 051126 | 4.154 | 0.210416 | $3.518 \times 10^{-5}$ | 0.059209 | 0.065653 | 0.020549 | N.D. | No | N.C. |
| Methanobacteriota archaeon | 2 | 051126 | 4.154 | 0.204014 | $3.411 \times 10^{-5}$ | 0.062680 | 0.065653 | 0.024020 | N.D. | No | N.C. |
| Methanobacteriota archaeon | 3 | 051126 | 4.154 | 0.207231 | $3.464 \times 10^{-5}$ | 0.058399 | 0.065653 | 0.019739 | N.D. | No | N.C. |
| GEN_4 | 1 | 042026 | 1.472 | 0.013188 | $6.221 \times 10^{-6}$ | 0.027448 | N.A. | N.C. | N.D. | N.C. | N.C. |
| GEN_4 | 2 | 042026 | 1.472 | 0.026232 | $1.238 \times 10^{-5}$ | 0.028016 | N.A. | N.C. | N.D. | N.C. | N.C. |

| Display name | Replicate | Source plate | Protein (mg/mL) | 3PGA (mM) | Apparent 3PGA specific activity (μmol $mg^{-1}$ $min^{-1}$) | Raw 2PG (mM) | 2PG threshold (mM) | Corrected 2PG (mM) | Apparent 2PG specific activity (μmol $mg^{-1}$ $min^{-1}$) | Passed raw-2PG threshold | Apparent specificity |
|---|---|---|---|---|---|---|---|---|---|---|---|
| GEN_4 | 3 | 042026 | 1.472 | 0.014016 | $6.612 \times 10^{-6}$ | 0.029245 | N.A. | N.C. | N.D. | N.C. | N.C. |
| Candidatus Gottesmaniibacteriota | 1 | 052026 | 2.036 | 0.026000 | $8.868 \times 10^{-6}$ | 0.032778 | 0.072594 | -0.016485 | N.D. | No | N.C. |
| Candidatus Gottesmaniibacteriota | 2 | 052026 | 2.036 | 0.021238 | $7.244 \times 10^{-6}$ | 0.037440 | 0.072594 | -0.011823 | N.D. | No | N.C. |
| Candidatus Gottesmaniibacteriota | 3 | 052026 | 2.036 | 0.024487 | $8.352 \times 10^{-6}$ | 0.047471 | 0.072594 | -0.001791 | N.D. | No | N.C. |
| GEN_10 | 1 | 051126 | 3.148 | 0.020822 | $4.593 \times 10^{-6}$ | 0.026507 | 0.065653 | -0.012154 | N.D. | No | N.C. |
| GEN_10 | 2 | 051126 | 3.148 | 0.019025 | $4.197 \times 10^{-6}$ | 0.023654 | 0.065653 | -0.015007 | N.D. | No | N.C. |
| GEN_10 | 3 | 051126 | 3.148 | 0.019642 | $4.333 \times 10^{-6}$ | 0.024078 | 0.065653 | -0.014583 | N.D. | No | N.C. |

Table S7. Primary and predefined sequence–structure–function test inventory. FDR correction was applied within the prespecified statistical families described in Methods. The single FDR-supported result was the signed basic-minus-acidic residue fraction versus soluble recovery ($q = 0.00980$). Absence of an FDR-supported association is not evidence of equivalence.

| Analysis family | Outcome(s) | Tests | n per test | q < 0.05 |
|---|---|---|---|---|
| Ligand-contact conservation | 3PGA, apparent 2PG, apparent specificity | 11 | 5–18 | 0 |
| Interfaces, packing, and active-site geometry | 3PGA, apparent 2PG, apparent specificity | 52 | 5–16 | 0 |
| Global sequence composition and solubility | Soluble recovery | 14 | 21 | 1 |
| Predicted structure and solubility | Soluble recovery | 13 | 21 | 0 |
| Selection metrics and solubility | Soluble recovery | 6 | 10–21 | 0 |
| Whole-protein similarity and solubility | Soluble recovery | 7 | 21 | 0 |
| Cavity geometry/electrostatics | 3PGA, apparent specificity, Tm | 24 | 9–17 | 0 |
| Total primary/predefined inventory | All predefined outcomes | 127 | 5–21 | 1 |

Table S8. Complementary sequence and whole-protein structural tests of residues, folds, dimers, interfaces, and static cavities. The three residue registers were analyzed independently to reduce dependence on primary-sequence alignment. These panel-specific screens test simple diagnostic explanations and do not establish universal residue necessity.

| **Analysis** | **Comparison** | **Result** | **Interpretation** |
|---|---|---|---|
| Sequence alignment | Strict active-conserved/nondetect-different residues | 0 | No single aligned residue separated all 23 3PGA-positive proteins from GEN_5 or 3C7 (Ca. Woesearchaeota). |
| 9RUB-guided structural register | Positions identical across all 23 positives | 34 | All 34 were also conserved in both nondetects. |
| Reference-independent US-align register | Strict substitutions; structurally unmapped positions | 0; 0 | The sequence result was unchanged in a whole-protein structural register. |
| 9RUB ligand-contact set | GEN_5 contact identity | 90% | Loss of the canonical contact set does not explain the GEN_5 nondetect. |
| Whole-protein US-align | Median pairwise TM-score, monomer; dimer | 0.921; 0.932 | Neither nondetect was a gross fold or dimer outlier. |
| Nearest whole-protein neighbors | GEN_5; 3C7 (Ca. Woesearchaeota) | GEN_8; 3D6 (A. boonei) | Each nondetect was closest to a 3PGA-positive protein. |
| Dimer interface | Strict active-conserved contacts lost in either nondetect | 0 | No diagnostic interface-contact loss was found. |
| Curated-coordinate CAVER | Primary; secondary FDR-supported relationships | 0/24; 0/30 | Static cavity descriptors did not explain 3PGA, apparent specificity, or Tm after correction. |

Table S9. GEN_10 data-collection and refinement statistics.

| | **GEN_10** |
|---|---|
| ***Data collection*** | |
| Space group | I 2 |
| Unit-Cell parameters (Å) | 64.498 83.784 73.944 90 104.96 90 |
| Resolution range (Å) | 35.72 - 1.40 |
| Total reflections | 253454 (12621) |
| Unique reflections | 73362 (3676) |
| $R_{merge}$ (%) | 0.066 (1.015) |
| $I/\sigma I$ | 9.8 (1.0) |
| Wilson B-factor | 13.3 |
| Completeness (%) | 98.4 (99.7) |
| Redundancy | 3.5 (3.4) |
| $CC_{1/2}$ | 0.998 (0.328) |
| ***Refinement*** | |
| Resolution range (Å) | 35.72 - 1.40 (1.44 - 1.40) |
| Reflections used in refinement | 73362 (3676) |
| Reflections used for $R_{free}$ | 1835 (134) |
| $R_{work}$ | 0.1492 (0.3022) |
| $R_{free}$ | 0.1697 (0.3447) |
| No. atoms | |
| Proteins | 3505 |
| Ligands/ion | 16 |
| Water | 319 |
| RMS from ideal geometry | |
| Bond lengths (Å) | 0.008 |
| Bond angles (º) | 0.91 |
| Average B-factor | 18.96 |
| Macromolecules | 17.81 |
| Ligands | 45.89 |
| Solvent | 30.29 |
| Ramachandran favored (%) | 97.03 |
| Ramachandran allowed (%) | 2.51 |

| | |
|---|---|
| Ramachandran outliers (%) | 0.46 |
| Clashscore | 5.01 |

[#] Values in parentheses are for highest-resolution shell.

Table S10. GEN_10 sequence–structure mapping to 35 eligible rigid active-site positions in *R. rubrum* Rubisco (PDB 9RUB). Reference residues were within 8 Å of RuBP or $Mg^{2+}$ after flexible loop 6 and C-terminal lid residues were excluded. Robust outlier rejection retained 31 positions as structural inliers for the local fit. Identity is conserved at 28 of 35 eligible positions (80.0%) and at 26 of 31 fit inliers (83.9%). KCX denotes carbamylated lysine and is counted as conserved relative to lysine.

| 9RUB residue | GEN_10 residue | Conserved identity | Local-fit inlier |
|---|---|---|---|
| GLY:162 | GLY:161 | Yes | Yes |
| THR:163 | THR:162 | Yes | Yes |
| ILE:164 | ILE:163 | Yes | Yes |
| ILE:165 | ILE:164 | Yes | Yes |
| LYS:166 | LYS:165 | Yes | Yes |
| LYS:168 | LYS:167 | Yes | Yes |
| LYS:191 | KCX:191 | Yes | Yes |
| ASN:192 | ASN:192 | Yes | Yes |
| ASP:193 | ASP:193 | Yes | Yes |
| GLU:194 | GLU:194 | Yes | Yes |
| PRO:195 | PRO:195 | Yes | Yes |
| LEU:261 | LEU:259 | Yes | Yes |
| ASP:263 | ASP:261 | Yes | Yes |
| HIS:285 | HIS:283 | Yes | Yes |
| TYR:286 | PHE:284 | No | Yes |
| HIS:287 | HIS:285 | Yes | Yes |
| ARG:288 | ARG:286 | Yes | Yes |
| ALA:289 | ALA:287 | Yes | Yes |
| HIS:291 | HIS:289 | Yes | Yes |
| CYS:309 | SER:307 | No | Yes |
| PRO:365 | PRO:364 | Yes | Yes |
| ILE:366 | ILE:365 | Yes | Yes |
| ILE:367 | ALA:366 | No | Yes |
| SER:368 | SER:367 | Yes | Yes |
| GLY:369 | GLY:368 | Yes | Yes |
| GLY:370 | GLY:369 | Yes | No |
| ILE:389 | ILE:388 | Yes | Yes |
| LEU:390 | THR:389 | No | Yes |
| THR:391 | THR:390 | Yes | Yes |
| ALA:392 | MET:391 | No | Yes |
| GLY:393 | GLY:392 | Yes | Yes |
| GLY:394 | GLY:393 | Yes | Yes |
| GLY:395 | GLY:394 | Yes | No |
| ALA:396 | VAL:395 | No | No |
| PHE:397 | HIS:396 | No | No |

Table S11a. Sequence-novelty measurements for the 21 experimentally tested designs: search outcome, experimental status, and identity and coverage of the closest natural match. Nearest-natural identity is the identity to the closest sequence in the curated natural Rubisco collection and is a different measurement from the UniRef90 identity in Tables S1 and S4. Query and target coverage are the fractions of the design and of the natural sequence covered by the alignment; no coverage filter was applied. Distinct matches are the number of matches remaining after grouping at 90% identity. Percentages and fractions are shown to two decimal places; the underlying file sequence_novelty_analysis/data/anti_chimera_metrics.csv holds the unrounded values.

| Design | Search status | 3PGA active | Soluble | Length (aa) | Nearest-natural identity (%) | Closest match | Closest-match identity (%) | Query coverage | Target coverage | Distinct matches |
|---|---|---|---|---|---|---|---|---|---|---|
| GEN_1 | hit | False | False | 448 | 77.34 | KYC52615.1 | 73.00 | 0.97 | 0.96 | 24 |
| GEN_2 | hit | False | False | 427 | 68.31 | NYZ77702.1 | 60.10 | 1.00 | 0.93 | 42 |
| GEN_3 | hit | False | False | 416 | 61.61 | MBN2517830.1 | 59.80 | 1.00 | 0.90 | 44 |
| GEN_4 | hit | True | True | 453 | 73.53 | VDS11140.1 | 69.40 | 0.94 | 0.94 | 29 |
| GEN_5 | hit | False | True | 447 | 67.83 | VDS11140.1 | 67.50 | 0.98 | 0.96 | 35 |
| GEN_6 | hit | False | False | 426 | 65.98 | NYZ77702.1 | 65.70 | 1.00 | 0.93 | 43 |
| GEN_7 | hit | True | True | 443 | 77.50 | NYT03909.1 | 73.70 | 0.98 | 0.98 | 24 |
| GEN_8 | hit | True | True | 435 | 80.36 | VDS11196.1 | 73.50 | 0.99 | 0.96 | 32 |
| GEN_9 | hit | False | False | 431 | 75.37 | VDS11140.1 | 69.90 | 0.96 | 0.95 | 25 |
| GEN_10 | hit | True | True | 443 | 76.39 | MBB1543276.1 | 71.00 | 0.99 | 0.96 | 33 |
| GEN_11 | hit | True | True | 425 | 70.60 | MBD3310098.1 | 65.60 | 0.91 | 0.92 | 41 |
| GEN_12 | hit | False | False | 439 | 74.64 | MBI4895876.1 | 71.40 | 0.96 | 0.96 | 41 |
| GEN_13 | hit | False | False | 460 | 72.49 | MBS3159242.1 | 66.40 | 0.93 | 0.97 | 34 |
| GEN_14 | hit | False | False | 444 | 57.70 | NYZ77702.1 | 57.30 | 1.00 | 0.94 | 45 |
| GEN_15 | hit | False | False | 447 | 75.42 | RLG91113.1 | 66.60 | 0.98 | 0.98 | 31 |
| GEN_16 | hit | False | False | 427 | 70.12 | RLG20169.1 | 70.10 | 0.99 | 0.99 | 44 |
| GEN_17 | hit | False | False | 431 | 71.32 | RLG20169.1 | 69.70 | 1.00 | 1.00 | 45 |
| GEN_18 | hit | False | False | 343 | 77.04 | EKD48085.1 | 71.40 | 0.99 | 0.78 | 34 |
| GEN_19 | hit | False | False | 441 | 72.14 | VDD88882.1 | 64.00 | 0.96 | 0.96 | 42 |
| GEN_20 | hit | False | False | 415 | 70.00 | MBI4895876.1 | 67.30 | 1.00 | 0.96 | 45 |
| GEN_21 | hit | False | False | 431 | 73.40 | VDD88882.1 | 68.20 | 0.99 | 0.99 | 44 |

Table S11b. Block-level sequence-novelty measurements for the same 21 designs. Cumulative identity is the fraction of the design covered by identical residues contributed by the closest one, two, three, and five distinct matches. The longest exact match is the longest run of consecutive identical residues shared with any distinct match, whereas the longest run in the closest match is the quantity evaluated by the permutation test. The permutation mean is the average longest run expected when identical residues are rearranged at random, and the Holm-adjusted P value is reported for the five designs that produced quantifiable 3PGA. Values are shown to two decimal places and P values to four significant figures; the underlying file sequence_novelty_analysis/data/anti_chimera_metrics.csv holds the unrounded values.

| Design | Cum. identity top 1 | Cum. identity top 2 | Cum. identity top 3 | Cum. identity top 5 | Longest exact match (aa) | Max 30-aa window identity | Longest merged ≥90% span (aa) | Longest same-match run | Longest run in closest match (aa) | Perm. mean run | Perm. P | Holm-adjusted P | Holm signif. |
|---|---|---|---|---|---|---|---|---|---|---|---|---|---|
| GEN_1 | 0.76 | 0.77 | 0.81 | 0.83 | 42 | 1.00 | 93 | 83 | 39 | 19.94 | 0.0037 | — | False |
| GEN_2 | 0.62 | 0.86 | 0.89 | 0.92 | 37 | 1.00 | 85 | 121 | 37 | 11.54 | 9.999e-05 | — | False |
| GEN_3 | 0.63 | 0.71 | 0.74 | 0.80 | 38 | 1.00 | 75 | 62 | 22 | 11.66 | 0.005 | — | False |
| GEN_4 | 0.66 | 0.74 | 0.75 | 0.77 | 52 | 1.00 | 135 | 120 | 31 | 14.79 | 0.002 | 0.007999 | True |
| GEN_5 | 0.67 | 0.73 | 0.74 | 0.77 | 36 | 1.00 | 59 | 71 | 31 | 13.67 | 0.0004 | — | False |
| GEN_6 | 0.67 | 0.72 | 0.75 | 0.78 | 37 | 1.00 | 80 | 92 | 37 | 13.38 | 9.999e-05 | — | False |
| GEN_7 | 0.77 | 0.77 | 0.83 | 0.86 | 42 | 1.00 | 89 | 164 | 34 | 20.58 | 0.0158 | 0.0474 | True |
| GEN_8 | 0.78 | 0.84 | 0.85 | 0.89 | 41 | 1.00 | 103 | 66 | 29 | 20.51 | 0.06859 | 0.1372 | False |
| GEN_9 | 0.68 | 0.75 | 0.77 | 0.79 | 38 | 1.00 | 82 | 74 | 27 | 14.82 | 0.005399 | — | False |

| Design | Cum. identity top 1 | Cum. identity top 2 | Cum. identity top 3 | Cum. identity top 5 | Longest exact match (aa) | Max 30-aa window identity | Longest merged ≥90% span (aa) | Longest same-match run | Longest run in closest match (aa) | Perm. mean run | Perm. P | Holm-adjusted P | Holm signif. |
|---|---|---|---|---|---|---|---|---|---|---|---|---|---|
| GEN_10 | 0.75 | 0.80 | 0.81 | 0.88 | 50 | 1.00 | 109 | 126 | 50 | 18.41 | 9.999e-05 | 0.0005 | True |
| GEN_11 | 0.60 | 0.70 | 0.74 | 0.80 | 19 | 0.93 | 43 | 40 | 16 | 12.37 | 0.1229 | 0.1372 | False |
| GEN_12 | 0.68 | 0.75 | 0.77 | 0.79 | 34 | 1.00 | 81 | 81 | 29 | 15.44 | 0.0048 | — | False |
| GEN_13 | 0.66 | 0.72 | 0.75 | 0.80 | 37 | 1.00 | 92 | 43 | 33 | 15.18 | 0.0017 | — | False |
| GEN_14 | 0.59 | 0.70 | 0.74 | 0.78 | 43 | 1.00 | 67 | 111 | 43 | 10.55 | 9.999e-05 | — | False |
| GEN_15 | 0.66 | 0.69 | 0.72 | 0.76 | 26 | 0.97 | 67 | 89 | 25 | 13.40 | 0.005799 | — | False |
| GEN_16 | 0.70 | 0.77 | 0.80 | 0.84 | 28 | 0.97 | 64 | 65 | 16 | 14.68 | 0.3337 | — | False |
| GEN_17 | 0.70 | 0.78 | 0.81 | 0.86 | 32 | 1.00 | 58 | 41 | 20 | 14.53 | 0.07369 | — | False |
| GEN_18 | 0.71 | 0.83 | 0.86 | 0.88 | 22 | 0.97 | 67 | 61 | 22 | 14.75 | 0.0466 | — | False |
| GEN_19 | 0.61 | 0.70 | 0.76 | 0.80 | 29 | 0.97 | 101 | 82 | 26 | 12.05 | 0.0012 | — | False |
| GEN_20 | 0.67 | 0.74 | 0.77 | 0.81 | 28 | 0.97 | 72 | 88 | 22 | 13.47 | 0.0193 | — | False |
| GEN_21 | 0.68 | 0.75 | 0.78 | 0.81 | 36 | 1.00 | 48 | 86 | 25 | 13.86 | 0.005999 | — | False |

**Supplementary sequences:**

>GEN_1
KVKSALVRVKKYLNERNIDYINLSLPDPKNGEYMMAVFHLVPGAELNMLQAAAEVAA
ESSTGTNFLVQTETPFSKEMNALVYQLDSDKNLVWMAYPWRLSDRDGNVQNILTYIIG
NVLGMKEISALKLLDVWFPPDMLQQYDGPSYTLDDMRRYLNVFGRPILGTIIKPKIGLTA
LDYAQVCYDFWVGGGDFVKNDEPQANQDFCPFNKMVKLVVKAMDKAAKATGNKKV
HSFNVTSSDFDTMIKRCTMVKKAGFEKGSYAFLIDGMTAGWMAVQTLRRRYPDVFVH
FHRASHGAFTRPENPIGFSVLVLSKFARLAGASGIHTGTAGVGKMKGKPKEDISAANNIL
KFVAEGHFFEQEWAKINKECAIASGGLNPLLLKPFIDTIGNADFITTMGAGCHAHPKGTR
AGAIALVQACEQWKKGITLKEAGKDYPELSAAIKFWGEKFV
>GEN_2
MAYGGYSYIDKKYKPDKDNDFIVLFWAKGSMPIEKIAEAIAAESSVGSWTKLKTMNDF
VWKNYRARVFKIVKVTKKSGFIKIAYPLEHFDLKNIPQFQASVLGNIFGLKELKELYVFD
ISFPKKYQKQFKGPKFGIKGIRKYLGTKKSRRPHVGTIVKPKVGLTPKEWANVAYEAWK
GGLDLVKDDENLVDQDFCKWKDRLHEVVKAVNKAEKETGEKKGHTLNISANITAMM
DRMDLVKELGNKYVMMDILTLGWSALQTARNYTKLPIHAHRAGHAMFDRNPNHGMS
MEVIAQFARMIGVDTLHIGTAYGKMSGGKKEVLHIEKEIENKFTKKTKENLSQKWYGIK
PVFAVASGGVYPGMVDKIIKFMGKDIIIQAGGGIHGHPDGTKAGAKAMRQAVDAYMK
KIPLKKYAKSHKELKKALKKWK
>GEN_3
MAYGGYSYLDPKYSPDENNDVLVLFWAKGSMPIEKIAEAIAAESSVGSWTKLKTMNDF
VWKHYRARVFKIIKVTKNSGFIWIAYPLEHFDVKNIPQFQASVLGNIFGLKELEELYVFDI
SFPKRYQKQFKGPKFGIKGIRKYLGTKKSRRPHVGTIVKPKVGLTPKEWANVAYQAWA
GGLDLVKDDENLVDQDFCRWKDRLHEVVKAIEKAEKETGEKKLYAANISASADEMLK
RAEFVKENGGNCAMIDIMTVGFSGLQMLRKHRLGLILHGHRAMHSAFTRNPKHGISML
VLAKLCRLAGIDQLHTGTVVGKMEGGAEEVKAINEFLKSKWGKIKPVMPIASGGLHPG
LVPALYKILGNDVIINFGGGIHGHPDGTLSGAKAAYQAVEAVVKGISLKEYAKTHKELA
KALEKWG
>GEN_4
MTKKKEWYEEEYWKKVFTATPDEIDPDEYIIATYYVKPRGGLSILGAAANIAAEQSTGT
WTKVPGETDEVRRKHVGKVIGVYEAPYYEWGIPDNITERQYILQIAFPWRNFGNQFSML
LTTTVGNISMAGDIKLLDLYLPKKYLEKFDGPSYTLEDMRKYLGVYDRPILGTIIKPKIGL
SPKEYADVCYDFWVGGGDFVKNDEPQADQDFCPFEKMVEDVRLAMDRAVKETGKKK
VHSFNVSASDFDTMIKRCELVKKAGFEKGSYAFLIDGITAGWMAVQTLRRKYPDVFIHF
HRASHGAYTRPENPFGFSVLVLSKFARLAGASGIHTGTAGVGKMKGSPEEDIIAAHNILN
LKGEGYFFDQSWGSIKPCCPIASGGLNPVLLKPFIDVVGTTDFITTMGAGCHAHPDGTKA
GATALVQACEAYKKGISIEEYAKDHKELAKAIEFYKKKK
>GEN_5
MDQSNRYADLSLKEDDLIAGGKHLLVAYKMKPAEGYGFLATAAHIAAESSTGTNVEVC
TTDDFTKGVDCLVYEIDKTKFDDIYKIMKAKASIAYISFPWRLFDRGGNVQNILTYIAGNI
FGMGELKALKALDCWFPKEMLEHYDGPSTTIDDMRKYLNVYNRPILGTIIKPKVGLSSK
EYADVCYNFWVGGGDFVKNDEPQADQDFCPFKDMVDEVRLAMDRAEKETGKKKIHS
FNISAADFDTMIKRCEYVRNVMKPGSYAFLIDGLTAGWMAVQTLRRRYPDVFIHFHRA
GHGAFTRPENPIGFSVLVLSKFARLAGASGIHTGTAGVGKMKGSPEEDVTAAHNILSIKG
EGYFFEQDWKKIKPCCPIASGGLNPTLLKPFIDVIGTTDFITTMGAGCHAHPDGTRAGAK
ALVQACEAYIQGIDIEKYAKTHKELAKAIEKFGKK

>GEN_6
MAYGGYSYIDKKYVPDSKNDFIVLFWAKGSMPIEKIAEAIAAESSVGSWTKLKTMNDF
VWKNYRARVFKIIKVTKNSGFIWIAYPLEHFDIKNIPQFQASVLGNIFGLKELKELYIFDIS
FPKRYQKQFSGPKYGIEGIRKLLGTKKSRRPHVGTIVKPKVGLTPKEWANVAYEAYSGG
LDLVKDDENLVDQDFCRWKERLHEVVKAIEKAEKETGEKKVYAANISDRNSRMVERV
DYLNSLGLKKNMVDIMTMGWSAVQELRNLNLKVILHRHRAMHAAFTRNPKHGISMLV
IAKLARLAGVDQLHTGTVVGKMEGGKEEVLAINEFLKSDFGHYHILRENWSKIKPTLPIA
SGGLHPGLLPKLYDILGENIIINFGAGIHGHPDGTYAGAKACVQAVEAIMKKIPLKKYAK
THKELAKALEHWK
>GEN_7
MKDDFSSLSAKEKDYINLDLPDVRKGEYLLAVFHLIPGGKLNILQAAAEIAAESSTGTNF
LVKTETEFSKEMNALVYKLDEKNFLAWIAYPWRLFDRGGNVQNILTYIAGNVFGMKEL
SALKLLDVWFPPSMLEQYDGPSYTLSDMRKYLGVYDRPILGTIIKPKIGLKPKEFADVCY
DFWVGGGDFVKNDEPQADQDFCPFEEMVDDVRLAMDKAEKETGKKKVHSFNVSASD
FDTMIKRAELVRKAGFEKGSYAFLIDGITAGWMAVQTLRRKYPDVFIHFHRAGHGAYT
RPENPIGFSVLVLSKFARLAGASGIHTGTAGIGKMKGDEKDDIVAANIALDKIANGNYFT
QDWYGMKPCCPIASGGLNPTKLKPFIDAIGTTDFITTMGAGCHAHPDGTKAGATALVQA
CEAWKKGITIEEYAKDHKELARAIEYFKKGVV
>GEN_8
MDKDYKYLNLGLDPKKNGNYLLVVYHLVPKDKLNFLQAAAEIAAESSTGTNLKVGTA
TDFSKSLNALVYKVDEKKNLVWIAYPWRIFDRGGNIQNILTYVVGNILGMKEIKALKLL
DVWFPPEMLSKYDGPSYTLDDMRKYLGVKDRPILGTIIKPKIGLTAAEYAEVCYDFWVG
GGDFVKNDEPQADQDFCPFDKMVDHVKKAMDKAVKETGKKKVHSFNVSASDYDTMI
KRCELVKSAGFEKGSYAFLIDGITAGWMAVQTLRRKYPDVFIHFHRAAHGAFTRKENPF
GFSVLVLSKFARLAGASGIHTGTAGVGKMSGSPEEDITAAHQALMLESDGYFFNQTWG
TIKPMCPIASGGLNPVLLKPFIEIIGNTDFITTMGAGCHAHPGGTKAGATALVQACEAWK
QGITIEEYAKTHEELKVAIEYYNKKKK
>GEN_9
KIQDTEYDPKSHYLMCVFFIYPKKGYTLLDAASEVAAESSTGTNFAVDTETDFSKEMNA
LVYKVDEEKLLVWIAYPWRIFDRGGNVQNILTYLAGNVFGMSTLNACKLLDCWFPPAM
LEQYDGPSYTLDDMRKYLGVYDRPILGTILKPKIGLRPEEFADACYDFWVGGGDFVKN
DEPQADQDFCPFEKMVDNVRAAMDKAVKQTGKKKVHSFNVSASDYDTMIERAKMISK
AGNEKGYYSFLIDGTTAGMMAVQTLRRKYPDVFIHFHRASHGAYTRPENPIGYSVLVLS
KFARLAGASGIHTGTAGVGKMAGSAEEDVAVANVILDDKAKGYFFDQDWKKIEPCSSI
ASGGLNPTKIHREVALVGDVIVITAGGQHGHPQGTHKGAKAAMDAIEAWKQGIPLLEY
AKSHKELKGAWEKFFSGSGREYRK
>GEN_10
MSEAKIIEALNRHQSAYVNLNLKNPRNGQYMLAAFHLIPGGKLNMLQAAAEVAAESST
GTNFMVETETPFSKEMNALVYKVDEEKELVWIAYPWRLFDRGGNVQNILTYIAGNVFG
MKEVKALKLLDVWFPPEMLEKYDGPSYTLSDMRNYLGVKDRPILGTIIKPKMGLTPKE
YADVCYDFWVGGGDFVKNDEPQADQDFCPFEEMVDEIRMAMDRAEQETGKTKVHSF
NVSASAYETMIRRADYVKKIMKPGSYAYLVDGLTAGWMAVQTLRRRYPDVFIHFHRA
SHGAYTRPENPFGFSVLVLSKFARLAGASGIHTGTAGVGKMKGDPEEDIKAASAILDLK
GEGYFFDQSWHSIKEMNPIASGGLNPTLLKPFIDIVGTVDFITTMGGGVHSHPGGTRKGA
MALVQACEAWKKGITIEEYAKDHEELRQAIEFYKRNK
>GEN_11

MTELSATRLRLPASLDPRRFAVATAGVDLPLDALATRLAAESSVGTWTDVATLSPALAR
RLRPHVLEVDADRGTVRIAYPSELFEAGNMPQVLSSVAGNVFGMKAVDGLRLLDIEFPD
SLVRAHPGPRFGIDGVRRLLGVEDRPLVGTIVKPKVGLDPERHAEVAYEAWRGGCDLV
KDDENLTDQDFNPFEKRVERTLKARDRAEEETGERKAYMPNVTAETEEMLRRAEFVKE
HGGEYVMVDILTVGWSALQTLREADLGLVIHAHRAMHAAFTRNPRHGISMLVVAKLA
RLAGVDQLHIGTAVGKMEGGREEVLEIEEEIEERFVRPARPAHVLEQDWFGIKPVFAVCS
GGLHPGHVPAVIERMGRDIIIQAGGGIHGHPDGTEAGARAMRQAVEAALRGVPLDEYA
RDHKELAKALKKWGRR
>GEN_12
MSAKKLSLPRFSVKEILDVESDLAVLYKIKPASGLSIEDAAGGVAAESSTGTWTTLYKW
CDKDRVKKLSAKAYSFIDLNDGSWIVKIAYPVELFEIGSIPNFLASIAGNIFGMKRVK
NLRLEDIYFPRKFIEKFKGPNYGRDGIRKLLKVKDRPLVGTVPKPKIGYSAKEHAKI
AYETWVGGFDLAKDDENLTSQKFNKFEERVRLMAKMRDKAEKETGERKDALINIT
AETKEMEKRAKILHDYGWRYAMIDVVVSGTSAVQTLRETLGDYGMAIHAHRAMH
ASFDRNPKHGITMSFLAKLMRMIGVDQIHSGTAVGKLVGGREEVQAIADVLREKK
TKPKEHFLLDQDWHNIKPAFPVSSGGLHPGLVPDIIDILGEDCIILVSGGIHGHPKGT
RAGAKAAMQAIEAYNKKIPLKEYAKTNIALAQALEKWGFMKPV
>GEN_13
MDEKDKYKMLYLKLDDKNIYEGNFYLAVYKMKPKEGLDLLGVASEIAAESSTGTNLK
VCTATPFSRDMDALVYEIDEENGIAYIAYPWRIFDRGGNVQNILTFVVGNVLGMGN
IEACKALDCWFPKEMLEHYDGPSTTIHDLKEYLGVKNRPVLGTIIKPKIGLSAKEYA
KVCYDFWIGGGDFVKNDEPQADQDFAPFEKMVDEVLESMDRAMSETGKNKVHSF
NVSASDFDTMIKRCELVRSVMKKGSYAFLVDGLTAGWMAVQTLRRRYPDVFIHFH
RAGHGAFTRPENPFGFSVLVLSKFARLAGASGIHTGTAGVGKMKGSPEEDIMAAN
NILSMKSEGYFFEQDWKTIRKCCPIASGGLNPTLLKPFIDLIGSTDFITTMGAGCHAH
PKGTKKGATALVQACEAYKKKISIKEYSKDHPELAEAIKFFSKKKEYVLPKSNVIPL
DSDKKLARPMV
>GEN_14
MVYGGYSYIDPKYEPKKDDFIVLLWAKGSEPIEKIAEAIAAESSVGSWTKLKTMNDFVW
ENLRARVFKIIKVTKKSGFIKIAYPLEHFDAKNISQFQASVLGNIFGLKELKELYAFDI
SFPKRYQKQFPGPLHGLEGIRKIIGTERSRRPHVGTIVKPKVGLTPKEWANVAYQA
WAGGLDLVKDDENLVDQDFCPWKDRFLKVLEAVNKAEEETGEKKLYAPNITAPA
DVMLKRAEFVKKNGGRYIMIDVVTTGFSALQMIRKYNPGLAIHAHRAMHAFMTR
ESGPGISGEGKIYNFSMSMIVIAKIMRLLGVDSLHIGSAKTKMHDSSEEEIIAIAEALR
GEMTPDKNLKMNQKWFGMKSVWPVASGGLHPGIVDKIVDVMGNDIFIQAGGGV
MGHPDGPRAGAKALRQAIEAYIKGISLKDYAKNHSELKQALEKWGYIRPV
>GEN_15
MVKKEWFEEEYWKKVFTATPDEIDPEKYIIATYYIKPKLGLSIKEAGASVAAEESIGTWT
KVYSGKNSGIKMAEKLRAVAYDLDLKNHMFKIAYKKELFEKGNMSGILAGIAGNI
DSMKMLKAFRLLDVRFPRKLIESFPGPRFGIDGIRKILNLKKRPITGTVPKPKVGYSP
EECGNLAYELLSGGMDYIKDDENLTSPSFCRFEKRAEKIMKAIEKAEKETGEKKVW
FANITSDIREMEKRMKLVADYGNPYVMVDVVIAGWSVLNYIRELAEDYGLAIHAH
RAMHAAFTRNPYHGISMFTLAKLYRVIGVDQLHIGTPEVGKLEAKTLDVIEYAKIL
RSNVYIPEENDIFHIKQEFHHIKPAMPVSSGGLHPGNIAKVVDILGEDIVLQVGGGVL
GHPDGPRSGARAVRQALEAIMKGIPLEEYAKTHRELAKALEKWGYVKPI
>GEN_16

MNYKEFIDLNYEPSESDLVVMFYFEPADGISKEEAIGRIASESSTGTWTTLHELPDRIKRL
KAKAYEFDGNYVKIAYPLDLWEPGNMAQLLSGIAGNIFGMKAVKNLRLIDVDLPD
KYIRGFKGPHYGREVKKKLLKIKKRPLLGTIPKPKVGFTAKEHADIAYQIWKGGIDL
IKDDENLTNQNFNRFEERVKKLAKVRDKAENETGDRKGALINITAETNEMIKRAKI
LHNLGWEYAMIDVVVTGTSALQTLRETLGDYGMAIHAHRAMHAAFDRNPKHGIT
MYFLAKLMRLIGVDQIHTGTAVGKLVGSKEEVLSIANTLRERKTKEKIRMILEQDW
YKIKPAFPVASGGLHPGLVPEIMKILGKDIVIQLGGGIHGHPKGTYSGAMAARQAV
DAYMEGISLEEKAKTSRELAQALEKWGYTKPK
>GEN_17
MAGKEWYHEFIDKSYKPKSTDLVVLFRFKVPRGMSVKEAAGRIASESSTGTWTTLYSLP
PRMKKLQATAFEIKGNYVKIAYPLDLWEPGNAPQLLSGIAGNIFGMKALNNLRLVD
IKLPKSYIKSFKGPSLGIKGIRKFMKIKKRPLTATVPKPKIGFSSKEHAKIGFETWMG
GFDLVKDDENLTSQKFNRFEERVKKMAKMRDKAEKETGERKSALINITAETKEMI
KRAKILKDLGWEYAMIDVVVTGTSALQTLREVLGDYGMAIHAHRAMHAAFDRNL
KHGITMATLAKLMRLIGVDQLHVGTAVGKLVSPREEVLNVSDALREKKVKPIPNLL
LDQNWYGIKDVLPVASGGLHPGLIDKIIDILGEDVVIQLGGGIHGHPKGTYSGAKAV
RQAIDAYKKGISLEEKAKNCKELKEALEKWGHEKPI
>GEN_18
DWCMDNIMVNISANVMGMKRVQGLRLEDVYMPPAFIKKFDGPSYTIEDMKKYLGIKN
RPILGTIVKPKLGLKPKEFADVCYNFWKGGGDFVKFDEPQADQNFCPFKEAVDEVL
KKMNKVKKETGKNKVMSINISAADFMTMQKRAEYVLKKMKKGSYAFLIDGLTAG
WTAVQTARRMYPDVFIHFHRAGHGAFTRPENAFGFSVPILTKFGRLAGASGIHTGT
AGIGKMSGSPEEDVMAAHNILDKKASGYFFDQDWKNIKPCCPIASGGLNPTKLHPF
IEAMGNTDFITTMGAGCHAHPDGTKAGAMALVQACEAYKQGVDISEYAKNHREL
ARAIEFYKKK
>GEN_19
MAVQLNYLAPPRWQPKNPDKNYLITYLDIELDKRANDNDFKDALASAAAESSRGTWTE
VYAGKGSGIKKADKKRAIAFDLDPKNHMFKIAYKQDLFEPGNMSGLLAGIAGNIDS
MKMLKAFRLIDVRFPEKLIRAFPGPKFGIKGIRKILKKKNRPLTGTVPKPKIGYTAKE
HAEVAYQTWLGGFDLTKDDENLTSMSFNNFYERVRLMTKLRDKAEQETGEVKDA
LINITAETKEMEKRAKILHNHGWRYAMIDVVVSGTSSVQTLRETLGDYGMAIHAH
RAMHASFDRNLKHGITMSFLAKIMRLIGVDQIHTGTAVGKLVGSKHEVLNIADTLR
ERHVKPYKNILLDQDWYSIKPAFPVSSGGLHPGLVPDILDILGKDIVIQVGGGVLGH
PKGARAGAMAVRQALEAKMKGISLDEYAKTHKELKQALKKWGFVKPM
>GEN_20
MSDIIATYYFHPKEDTTPEWAAQAIAEEQTTGTWTDISTTTSYVHYLDGRVVSIEKSTGY
YKVSIPLPQFDFPSILQLLAGVAGNIFGMKAVDNLRLVDVQFPEDYVNYYKGPKFGI
EGVRKYMKIKKRPLTGAVPKPKIGFSAKEHADIAFQTWMGGFDLVKDDENLTSMK
FNKFDDRVKLMTKLRDKAEKKTGDKKDALINITAETKEMEKRAKILHDYGWRYA
MIDVVVSGTSSVQTLRETLGDYGMAIHAHRAMHASFDRNLKHGMTMSALAKLMR
LIGVDQIHTGTAVGKLVGDREEVMSVANILREKKTKAFPKLLLEQDWGSIRTAFPV
ASGGLHPGLVPDVLDILGKDCIIQLGGGIHGHPKGTYYGAKAARQAIEAYMNKIPL
DEYAKTNKELAQALKKWGFMKPV
>GEN_21
MKKLEWYKEFVDLSYAPKKDDLILKYYVEPAKGISIKEAAGRVASESSAGTWTTLYTLP
ARMRKLQATAFEIDKNYIKVAYPLDLWEKGNAPQLLSGIAGNIFGMKALKNLRLID

IQFPKKYINSFKGPSHGIKGVRDILKVKDRPTTGAVPKPKIGFSAKEHADVAYETW
MGGFDLIKDDENLSSTDFNKFYKRVKLMTKMRDKAEKKTGEKKDALINITSETKE
MEKRAKLLHDYGWRYAMIDVVVTGFAAVQTVRNNLGDYGMALHAHRAMHALL
DRNPKHGITMSMLAKLMRMIGVDQIHVGTVVGKLVGGKDEVLNIRDAIVKENIKG
NKMSFLDQNWYNIKPAVPVSSGGLHPGLIPDIMQIMGKDCIIQVGGGIMGHPKGAY
AGAKAVRQALESKIKGISLKDYAKSNVELKEALDKWGQSKPV
